# Timing of transient darkness shapes carbon–nitrogen metabolism and sugar signaling in sugarcane

**DOI:** 10.64898/2026.09.01.748396

**Authors:** Gabriel M. Leal, Hellen O. de Oliveira, Felipe Farineli, Arthur Vanni-Lopes, William V. de M. Mira, Amanda F. Macedo, Grayce Hellen Romim, Carlos T. Hotta, Eny I. S. Floh, Bruno V. Navarro, Marcos S. Buckeridge

**Author notes:** corresponding author: Marcos Silveira Buckeridge. Gabriel Marques Leal, Hellen Oliveira de Oliveira, Felipe Farineli, Arthur Vanni-Lopes, William Vinícius de Mello Mira, Amanda Ferreira Macedo, Grayce Hellen Romim, Carlos Takeshi Hotta, Eny Iochevet Segal Floh, Bruno Viana Navarro, Marcos Silveira Buckeridge.

## Abstract

Fluctuating light is common in field environments. Yet, the mechanisms by which C4 crops coordinate carbon and nitrogen metabolism during short-term carbon deprivation remain poorly understood. Here, we imposed transient darkness at different phases of the diel cycle to assess how the timing of light loss affects photosynthesis, carbohydrate turnover, amino acid dynamics, and sugar-sensing pathways in commercial sugarcane leaves. Early-day darkness significantly impaired photosynthetic induction and revealed a temporal disconnect between stomatal and metabolic limitations, whereas midday and late-day treatments caused temporary, time-specific disruptions in carbon assimilation. These shifts altered the balance between sucrose preservation and catabolic mobilization, leading to treatment-dependent changes in starch reserves and free amino acids. Core circadian components largely maintained their phase relationships, but their amplitudes varied across treatments, consistent with partial decoupling from carbon status. Darkness also reorganized energy signaling, with SnRK1 and DIN6 responses associated with greater declines in sucrose. Notably, trehalose-pathway transcripts showed marked changes in network connectivity, with ScTPSIIG consistently emerging as a highly connected candidate associated with photosynthetic performance, water-use traits, sugar sensing, and amino acid metabolism. Overall, these results indicate that the timing of carbon limitation and residual sucrose availability shape distinct metabolic responses, while trehalose metabolism provides a candidate regulatory layer coordinating carbon–nitrogen adjustment during the diel cycle, highlighting class II TPS proteins as targets for functional investigation of metabolic resilience in sugarcane.

**HIGHLIGHTS:** Transient darkness disrupted sugarcane physiology in a time-dependent way.

Residual sucrose availability distinguishes buffered adjustment from stronger carbon starvation.

Network remodeling shows metabolite-centered coordination under light deprivation

Class II TPS isoforms emerge as candidate hubs in integrated metabolic responses.

## Introduction

Plant growth and productivity depend on the precise coordination of carbon assimilation, storage, and utilization throughout the diel cycle (Kölling et al., 2015; Smith & Zeeman, 2020; Liu et al., 2025). Beyond serving as metabolic substrates, soluble sugars also function as key signaling molecules that integrate environmental cues with cellular energy status, thereby regulating gene expression, enzyme activities, and developmental processes (Rolland et al., 2006; Li & Sheen, 2016). Diel regulation of carbohydrate metabolism, driven by light–dark transitions and the circadian clock, ensures a balanced supply of carbon skeletons to support respiration, growth, and biosynthesis at night (Smith & Stitt, 2007; Stitt & Zeeman, 2012). During the light period, photosynthesis promotes sucrose export and the buildup of temporary carbon reserves, while in darkness, these reserves are mobilized in a controlled manner to avoid nocturnal carbon starvation (Flis et al., 2019; Viana et al., 2021). In this context, sugar sensors such as Target of Rapamycin complex 1 (TORC1) and Sucrose non-fermenting-1 related kinase 1 (SnRK1) act antagonistically to regulate the switch between anabolic and catabolic states, whereas trehalose-6-phosphate (Tre6P) serves as an indicator of sucrose status, influencing both photosynthetic efficiency and sink–source relationships (Dobrenel et al., 2016; Figueroa & Lunn, 2016). The interplay between these signaling networks and the circadian oscillator provides a temporal framework for optimizing energy distribution under changing light conditions, linking carbon availability to time-dependent transcriptional and metabolic responses (Haydon *et al*., 2017; Viana *et al*., 2021).

Sugarcane (*Saccharum* spp.), a C4 grass of great agricultural importance, exhibits strong daily rhythms in photosynthesis and carbohydrate distribution (De Souza et al., 2018; Dantas et al., 2021). As a crop with high sucrose storage capacity and limited temporary starch reserves in leaves, its yield is especially sensitive to daily changes in light and to the timing of carbon supply to source tissues (Ribeiro *et al*., 2017; De Souza *et al*., 2018). Field and greenhouse research have shown that sucrose, organic acids, amino acids, and other key metabolites fluctuate in concert over the daily cycle in sugarcane leaves, with sucrose often following a different phase than other soluble sugars, suggesting additional layers of internal regulation beyond immediate photosynthesis (Lobo et al., 2015; Oliveira et al., 2025). These dynamics indicate that disruptions in light patterns at specific times can not only reduce carbon gain but also selectively influence the balance between sucrose export to sinks and the maintenance of metabolic balance in source tissues (Kölling et al., 2015; Wu et al., 2025).

Despite these advances, the effects of transient carbon limitation on sugarcane metabolism remain poorly understood. Previous studies have primarily used continuous darkness, extended nights, or prolonged shading. These approaches have provided important insights into sustained low-energy responses, including autophagy, amino acid accumulation, and SnRK1-associated signaling (Brouquisse et al., 1998). However, these treatments do not capture the effects of brief interruptions in photosynthetic carbon supply during a normal diel cycle. The metabolic response may depend on the timing of the light reduction. Carbohydrate reserves, metabolic activity, and circadian phase differ substantially between the beginning and the end of the photoperiod (Kölling et al., 2015). It remains unclear how rapidly sugarcane adjusts its carbon and nitrogen metabolism to short-term changes in light availability. The contributions of sugar-signaling pathways and clock-associated regulation to these responses also remain unknown.

Addressing these gaps is especially important for sugarcane, which often experiences temporary reductions in incident radiation due to cloud cover, self-shading in dense canopies, and seasonal changes in daylength. In this context, rapid adjustments in carbon allocation, amino acid metabolism, and energy signaling may help sustain sugar metabolism under fluctuating light conditions. Here, we applied short periods of local darkness at different phases of the diel cycle to induce transient limitations in photosynthetic carbon supply in sugarcane leaves. We combined gas exchange measurements with quantification of non-structural carbohydrates and free amino acids. We also analyzed the expression of transcripts associated with TORC1, SnRK1, the Tre6P pathway, and the circadian clock. We aimed to determine how the timing of light interruption affects carbon and nitrogen metabolism and low-energy signaling. This integrated approach reveals time-dependent responses to transient darkness in a major C4 bioenergy crop. It also identifies candidate regulatory components that may contribute to short-term metabolic adjustment in sugarcane.

## Material and methods

### Plant material and environmental conditions

Vegetatively propagated sugarcane plants (*Saccharum* spp. cv. SP80-3280) were grown outdoors under natural light from March to May 2024, during autumn in the Southern Hemisphere. Plants were cultivated in 15 L pots filled with a commercial forest-based substrate (Tropstrato HT Hortaliças®, Vida Verde, Brazil). The experiment was conducted at the Laboratório de Fisiologia Ecológica de Plantas, Universidade de São Paulo, São Paulo, Brazil (23°33′58.6″ S, 46°43′49.8″ W). Pots were arranged in a randomized block design. The substrate was initially supplemented with NPK fertilizer at a 30:20:30 ratio, followed by biweekly applications of Hoagland and Arnon nutrient solution (Hoagland and Arnon, 1950).

The experiment spanned a single diel cycle, from dawn on May 23 (ZT0) to dawn on May 24 (ZT24). At the start, plants were 60 days old and experienced approximately 11 h of natural light followed by 13 h of darkness. All pots were irrigated one hour before dawn and again at ZT6 to minimize variation in soil water availability. Environmental conditions were recorded hourly at canopy height in three sectors of the experimental area, corresponding to the western edge, center, and eastern edge (Fig. S1). Air temperature, relative humidity, and atmospheric CO₂ concentration were measured with a multifunction air quality meter (Testo 435-4, Testo, Germany). Photosynthetically active radiation was measured with a full-spectrum quantum sensor (MQ-500, Apogee Instruments, USA).

### Experimental design and dark treatments

We established five treatments to determine how the timing of transient local darkness affects leaf physiology and metabolism. Darkness was imposed on the youngest fully expanded leaf, designated leaf +1, and sampled at seven *Zeitgeber times* (ZTs; hours after dawn) by covering the entire leaf blade with aluminum foil. This procedure prevented light exposure to leaf +1 while the remaining leaves and the rest of the plant were maintained under natural light. We applied treatments during periods associated with previously reported metabolic changes in sugarcane throughout the diel cycle (de Souza et al., 2018). The treatments included: control without darkening (Natural light), darkness from ZT0–ZT4 (Morning dark), ZT4–ZT8 (Noon dark), ZT8–ZT12 (Afternoon dark), and ZT0–ZT12 (Constant dark). All plants were then exposed to the natural night period, from approximately ZT12 to ZT24. Leaf gas exchange was measured at ZT1, ZT4, ZT8, ZT12, ZT16, ZT20, and ZT24. Immediately after each measurement, we collected leaf +1 for carbohydrate, amino acid, and gene expression analyses. Five independent plants were sampled for each combination of treatment and collection time, with each plant representing one biological replicate. Supplementary Fig. S2 provides a schematic overview of the experimental design, including the timing of darkness treatments, sampling points, and sample collection strategy.

### Leaf gas-exchange measurements

We monitored gas-exchange dynamics at seven sampling times (ZT1, ZT4, ZT8, ZT12, ZT16, ZT20, and ZT24) throughout a full diel cycle across all treatments. Net CO₂ assimilation rate (*A*), stomatal conductance (g_s_), transpiration rate (*E*), intercellular CO₂ concentration (C_i_), and leaf-to-air vapor pressure deficit (Vpd_L_) were measured on the middle portion of leaf +1 at each sampling point using a portable photosynthesis system (LI-6400/LI-6400XT; LI-COR Inc., USA). Leaf temperature (T_i_), photosynthetic photon flux density (PPFD), and relative humidity followed ambient conditions at each sampling time. During induced dark treatments, PPFD was set to zero. Reference CO₂ concentration and airflow rate were maintained at 400 µmol mol⁻¹ and 300 µmol s⁻¹, respectively.

### Non-structural carbohydrates quantification

Non-structural carbohydrates were measured in the same leaf +1 samples used for prior gas exchange measurements, following De Souza et al. (2013). Immediately after measuring net CO₂ assimilation, leaf blades were detached, flash-frozen in liquid nitrogen, lyophilized, and ground to a fine powder. We extracted 10 mg of dry weight in four aliquots with 1 mL of 80% (v/v) ethanol warmed to 80 °C. We concentrated the combined supernatants under vacuum, redissolved them in 1 mL of Milli-Q water, and then partitioned them with 0.5 mL of 99% chloroform. Soluble sugars (glucose, fructose, sucrose, and raffinose) were quantified in the aqueous phase by high-performance anion-exchange chromatography with pulsed amperometric detection (HPAEC/PAD) using a CarboPac PA1 column (Dionex ICS-3000; Thermo Fisher Scientific, USA) with isocratic elution of 100 mM NaOH.

For starch extraction and quantification, wash the pellets obtained after ethanol extraction with distilled water and dry them overnight at 60 °C. Pellets were sequentially treated with α-amylase (120 U mL⁻¹; *Bacillus licheniformis*) and amyloglucosidase (*Aspergillus niger*) (Megazyme®, Ireland). We measured the glucose produced using a glucose oxidase/peroxidase (GOD/POD) assay kit (Labtest®, Brazil) and read absorbance at 490 nm. We used a glucose standard curve (20–300 mg mL⁻¹) for calibration and estimated starch content, assuming starch accounted for 90% of the total glucose released (Amaral et al., 2007).

### Determination of free amino acids

The duplicate aqueous-phase aliquots used for soluble sugar quantification were also used for free amino acid extraction and analysis, following de Oliveira et al. (2018) with minor modifications. Amino acids were derivatized with o-phthalaldehyde and separated by high-performance liquid chromatography (HPLC; Shimadzu, Japan) on a C18 reversed-phase column (5 µm × 4.6 mm × 250 mm; Supelcosil LC-18, Sigma-Aldrich, USA). Fluorescence detection was performed at 250 nm excitation and 480 nm emission. Amino acids were identified and quantified using retention times and peak areas, compared with reference standards: aspartate (Asp), glutamate (Glu), asparagine (Asn), serine (Ser), glutamine co-eluted with histidine (Gln + His), glycine (Gly), arginine (Arg), threonine (Thr), alanine (Ala), tyrosine co-eluted with γ-aminobutyric acid (Tyr + GABA), methionine (Met), tryptophan (Trp), valine (Val), phenylalanine (Phe), isoleucine (Ile), leucine (Leu), ornithine (Orn), lysine (Lys), and citrulline (Cit).

### qRT-PCR analysis

Total RNA was extracted from 200 mg of the same leaf +1 samples used for gas-exchange measurements and metabolite analysis using TRIzol® reagent and the PureLink RNA Mini Kit (Thermo Fisher Scientific,hoa USA) according to the manufacturer’s instructions. We measured RNA concentration and purity spectrophotometrically (NanoDrop 2000; Thermo Fisher Scientific) and used only samples with A260/280 ratios between 1.8 and 2.2 and A260/230 ratios between 1.6 and 2.2 for further analyses. After DNase I treatment (Thermo Fisher Scientific), first-strand cDNA was synthesized from 1 µg of total RNA using the SuperScript III First-Strand Synthesis System (Thermo Fisher Scientific).

Quantitative real-time PCR (qRT-PCR) was performed using Power SYBR Green PCR Master Mix on a QuantStudio 6 Flex Real-Time PCR System (Applied Biosystems, USA). Each 14 µL reaction contained 1.4 µL of 1:10-diluted cDNA and 400–800 nM of each primer, depending on the target. All reactions were run in duplicate across three biological replicates per treatment, and no-template controls (NTCs) were included to verify reaction specificity and ensure no contamination.

The analyzed transcripts included targets for sugar sensing (*ScTOR* and *ScSnRK1α*), trehalose metabolism (*ScTPS I, ScTPS II, ScTPP,* and *ScTRE*), circadian regulation (*ScLHY, ScTOC1, ScPRR37,* and *ScPRR73*), and energy-stress responses (*ScDIN6* and *ScbZIP63*). Transcript sequences for trehalose pathway components were obtained from de Oliveira et al. (2022), and those for *ScTOR* and *ScSnRK1α* from de Oliveira et al. (2024). We adopted primers for circadian clock genes from Hotta et al. (2013) and Dantas et al. (2021). Table S1 provides the complete list of primer sequences. Relative expression levels were calculated using the 2^⁻ΔΔCt^ method (Livak & Schmittgen, 2001) after baseline correction with LinRegPCR. We normalized expression values to the geometric mean of *ScGAPDH*, *ScACT,* and *ScPGR*.

### Data preprocessing and network analyses

We identified outliers independently for each combination of sampling time, treatment, and variable using Tukey’s interquartile range (IQR) criterion (Q1 − 1.5 × IQR and Q3 + 1.5 × IQR). We imputed missing transcript expression values to match the sample size of other datasets (n = 5) using the *k*-nearest neighbors (KNN) algorithm (Troyanskaya *et al.,* 2001), implemented in scikit-learn (Pedregosa *et al.,* 2011), with k = 3 and uniform weighting. We used the resulting dataset, comprising 13 transcript expression variables and 38 physiological and metabolic parameters, for weighted network analysis.

We built treatment-specific networks using the BioNetStat package (Jardim *et al.,* 2019) and visualized them in Cytoscape 3.8.0 (Shannon *et al.,* 2003). We estimated pairwise associations among the 51 measured variables using Spearman’s rank correlation coefficients. Weighted networks were inferred using the absolute correlation coefficient, and edges were retained when |r| > 0.8. To interpret treatment-induced network rewiring, we performed differential network analyses comparing each darkness treatment with the Natural Light control. We evaluated network topology using four centrality metrics (degree, betweenness, closeness, and eigenvector centrality) and assessed statistical significance via permutation testing with 1,000 label permutations. We also evaluated treatment-specific changes in network organization using either the complete dataset (all variables) or seven functional subsets (all transcripts, sugar sensing targets, circadian clock targets, all metabolites, non-structural carbohydrates, amino acids, and gas exchange).

### Multivariate and correlation analysis

We performed Wilcoxon tests (*P* < 0.05) and generated heatmaps in R version 4.3.2 (R Core Team, 2023). Correlation heatmaps were generated using Spearman’s rank correlation coefficients, considering only associations with |r| > 0.8. To evaluate multivariate discrimination among *Zeitgeber times* within each darkness treatment, we performed partial least squares discriminant analysis (PLS-DA) and calculated variable importance in projection (VIP) scores using MetaboAnalyst 5.0 (Chong *et al.,* 2019). Before analysis, we log_10_-transformed all 51 variables, mean-centered them, and divided them by each variable’s standard deviation (Fig. S3). We evaluated model performance using five-fold cross-validation with the first five latent components (Fig. S4). We assessed model performance using classification accuracy, R², and Q², and evaluated statistical significance by permutation testing with 1,000 permutations (Fig. S5-6).

## Results

### The timing of transient local darkness influences diel carbon assimilation

To determine how the timing of transient darkness affects photosynthetic performance, we monitored leaf gas exchange throughout a complete diel cycle under Natural Light and during induced darkness treatments (Fig. 1). Under Natural Light, net CO₂ assimilation increased rapidly after dawn and peaked at ZT4, reaching 20.39 µmol CO₂ m⁻² s⁻¹. Assimilation then declined after ZT8 as incident irradiance decreased, consistent with the diel pattern previously reported for field-grown sugarcane (de Souza et al., 2018). Stomatal conductance followed a similar daytime pattern, whereas transpiration increased until ZT8, associated with a higher leaf-to-air vapor pressure deficit and lower relative humidity (Figs. 1 and S1). After ZT20, stomatal conductance and transpiration increased slightly under Natural Light, coinciding with higher nocturnal relative humidity.

**Fig. 1.**
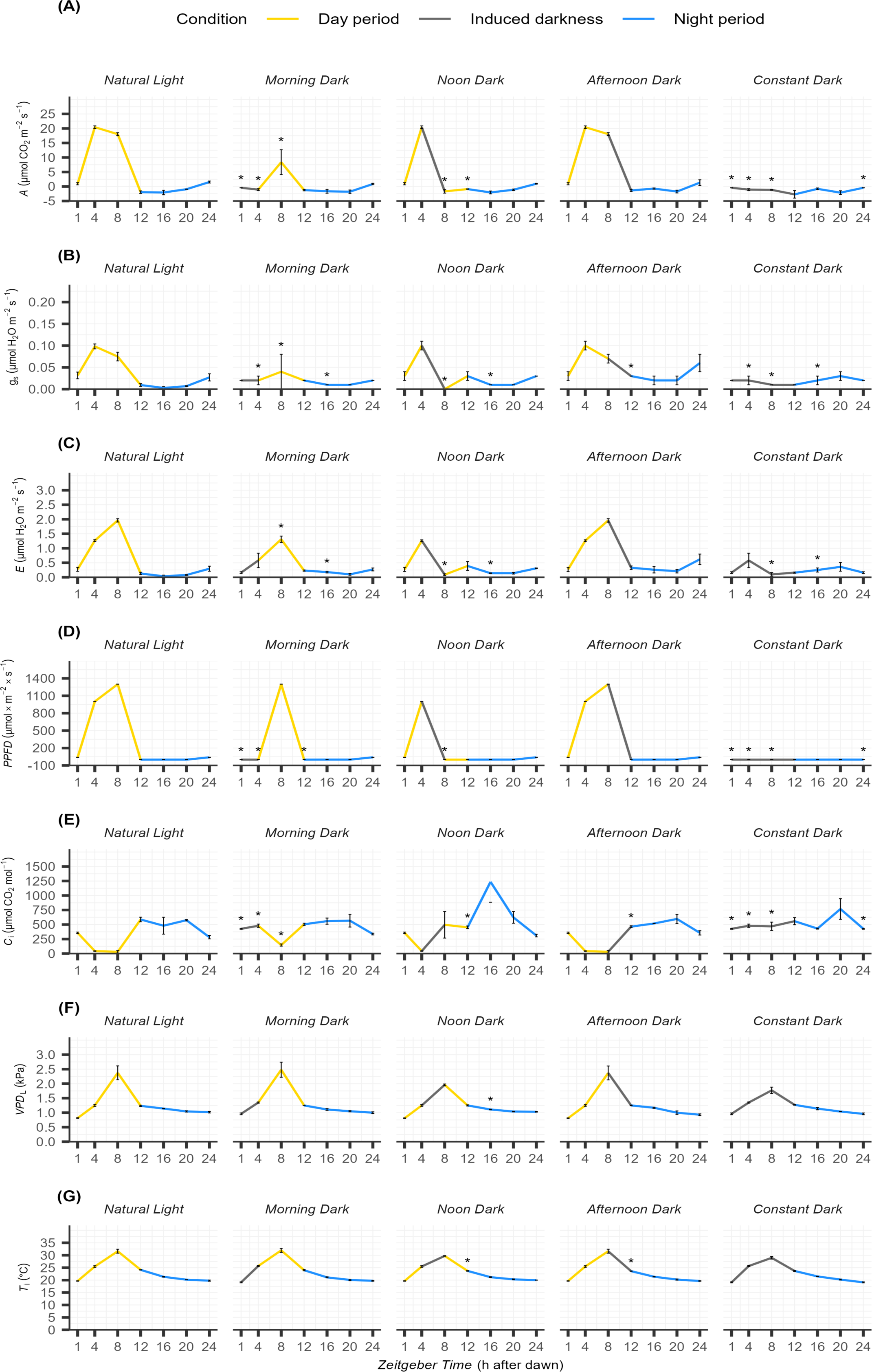
Diel variation in gas exchange and leaf microenvironment under different periods of induced darkness in sugarcane leaf +1. (A) Net CO₂ assimilation rate - *A*, (B) stomatal conductance - *g_s_*, (C) transpiration rate - *E*, (D) photosynthetic photon flux density - PPFD, (E) intercellular CO₂ concentration - C_i_, (F) leaf to air vapor pressure deficit - VPD_L_, and (G) leaf temperature - T_i_ were measured at seven *Zeitgeber times* (ZT1, ZT4, ZT8, ZT12, ZT16, ZT20, and ZT24). ZT indicates the number of hours elapsed since dawn. Treatments comprised Natural Light, Morning Dark from ZT0 to ZT4, Noon Dark from ZT4 to ZT8, Afternoon Dark from ZT8 to ZT11, and Constant Dark from ZT0 to ZT12. Yellow, black, and blue line segments represent measurements obtained during the natural light period, induced darkness, and natural night, respectively. Values are means ± SE of five biological replicates. Asterisks indicate significant differences between each induced darkness treatment and Natural Light at the same ZT according to the Wilcoxon test at P < 0.05.

In the Morning Dark treatment, net CO₂ assimilation remained near zero while the leaf was covered and recovered only partially after reillumination. At ZT8, approximately four hours after light exposure was restored, assimilation reached 8.37 µmol CO₂ m⁻² s⁻¹ and remained below the value measured under Natural Light. Stomatal conductance and transpiration also showed incomplete recovery during the remaining light period (Fig. 1). The effects of darkness applied later in the photoperiod differed from those observed in Morning Dark. In the Noon Dark treatment, net CO₂ assimilation remained lower than under Natural Light after reillumination at ZT8 and showed no clear recovery before the onset of the natural night. In Afternoon Dark, the values measured at the available sampling times did not differ significantly from those under Natural Light. However, the four-hour interval between measurements limited detection of short-term changes during the final portion of the photoperiod. In Constant Dark, net CO₂ assimilation remained near zero throughout the natural light period, confirming that the leaf covering effectively prevented photosynthetic carbon assimilation (Fig. 1A).

### Transient darkness alters the diel leaf partitioning of carbon between sucrose and starch

To examine how transient darkness affects leaf carbon balance, we measured concentrations of glucose, fructose, raffinose, sucrose, and starch throughout the diel cycle (Fig. 2). Under Natural Light, glucose and fructose followed similar profiles, with small daytime fluctuations and lower concentrations toward the end of the natural night (Fig. 2C and D). Raffinose also remained relatively stable, except for a transient decrease at ZT16, when its concentration reached 0.093 µg mg⁻¹ DW (Fig. 2E). Sucrose accumulated during the light period, peaked at ZT8 at approximately 30 µg mg⁻¹ DW, and then declined, remaining between 21 and 25 µg mg⁻¹ DW during the night (Fig. 2A). Starch showed the greatest diel variation, increasing sharply between ZT4 and ZT8 and reaching 46.76 µg mg⁻¹ DW. The starch pool then progressively decreased during the night and reached approximately one third of its daytime maximum by ZT24 (Fig. 2B).

**Fig. 2.**
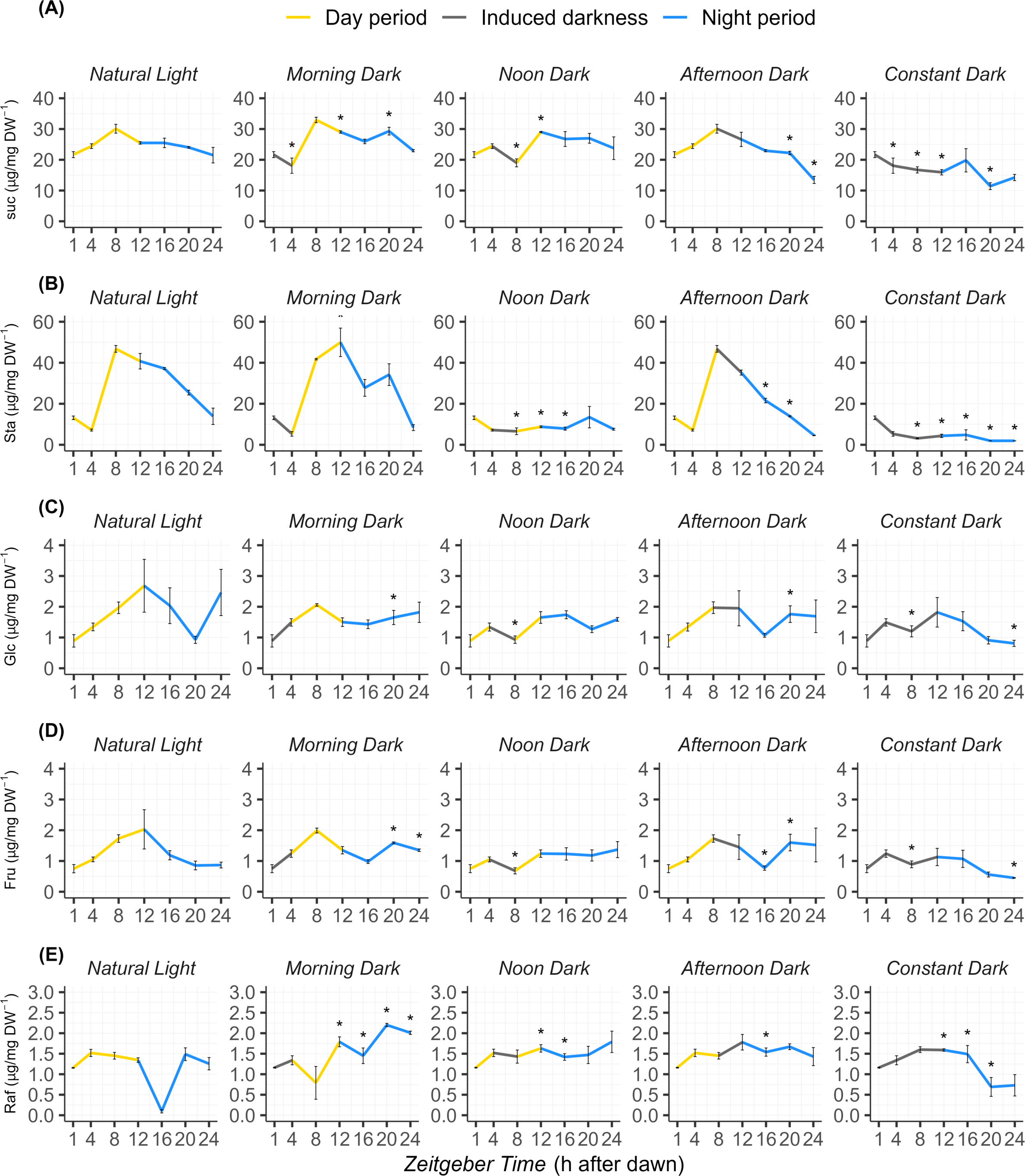
Diel variation in nonstructural carbohydrate concentrations in sugarcane leaf +1 under different periods of induced darkness. (A) Sucrose - Suc, (B) starch - Sta, (C) glucose - Glc, (D) fructose - Fru, and (E) raffinose - Raf were quantified at seven *Zeitgeber times* (ZT1, ZT4, ZT8, ZT12, ZT16, ZT20, and ZT24). ZT indicates the number of hours elapsed since dawn. Treatments comprised Natural Light, Morning Dark from ZT0 to ZT4, Noon Dark from ZT4 to ZT8, Afternoon Dark from ZT8 to ZT11, and Constant Dark from ZT0 to ZT12. Yellow, grey, and blue bars represent samples collected during the natural light period, induced darkness, and natural night, respectively. Grey shaded areas indicate the duration of induced darkness within each treatment. Carbohydrate concentrations are expressed as µg mg⁻¹ dry weight. Values are means ± SE of five biological replicates. Asterisks indicate significant differences between each induced darkness treatment and Natural Light at the same ZT according to the Wilcoxon test at P < 0.05.

The Morning Dark treatment altered both starch and soluble sugar profiles. After reillumination, starch accumulated to 49 µg mg⁻¹ DW at ZT8, the highest value observed across treatments. Its concentration then declined by approximately 80 to 90% from the daytime maximum to ZT24. A similarly pronounced nocturnal decline was observed in Afternoon Dark (Fig. 2B). In Morning Dark, sucrose remained relatively high after reillumination and throughout the subsequent night, while raffinose increased to between 1.45 and 2.20 µg mg⁻¹ DW. In contrast, sucrose progressively decreased in Afternoon Dark and reached 13.5 µg mg⁻¹ DW at ZT24, the lowest value observed for this treatment.

Noon Dark produced a distinct carbohydrate profile. Starch concentrations remained low after light exposure was restored at ZT8 and did not recover to the levels observed under Natural Light (Fig. 2B). Sucrose declined during the imposed darkness but increased after reillumination, along with a rise in raffinose. Despite these changes, sucrose concentrations remained between 23.76 and 29.04 µg mg⁻¹ DW for most of the cycle, within the range observed under Natural Light. In Constant Dark, starch remained low throughout the experiment and declined from 5.35 to 1.94 µg mg⁻¹ DW. Sucrose also decreased progressively, from 21.67 µg mg⁻¹ DW at the beginning of the cycle to 11.42 µg mg⁻¹ DW at ZT24 (Fig. 2A and B).

### The timing of transient darkness enhances free amino acid accumulation

To examine how transient darkness affects amino acid metabolism, we measured total and individual free amino acid concentrations throughout the diel cycle (Fig. 3). Under Natural Light, total free amino acid concentrations remained relatively stable, except for an increase at ZT8 that reached 18,033.73 nmol g⁻¹ DW. This increase was accompanied by higher concentrations of nine amino acid signals (Fig. S7). Morning Dark showed a similar diel profile, although the ZT8 increase was less pronounced and reached 15,263.77 nmol g⁻¹ DW. Despite this similarity in total concentration, the relative abundance of individual amino acids differed from Natural Light after the onset of darkness. Arg and the combined Gln+His signal increased, whereas Ala, Asp, Glu, and Ser decreased (Fig. S8). In Afternoon Dark, several amino acids accumulated predominantly toward the end of the night, particularly between ZT20 and ZT24 (Fig. S10).

**Fig. 3.**
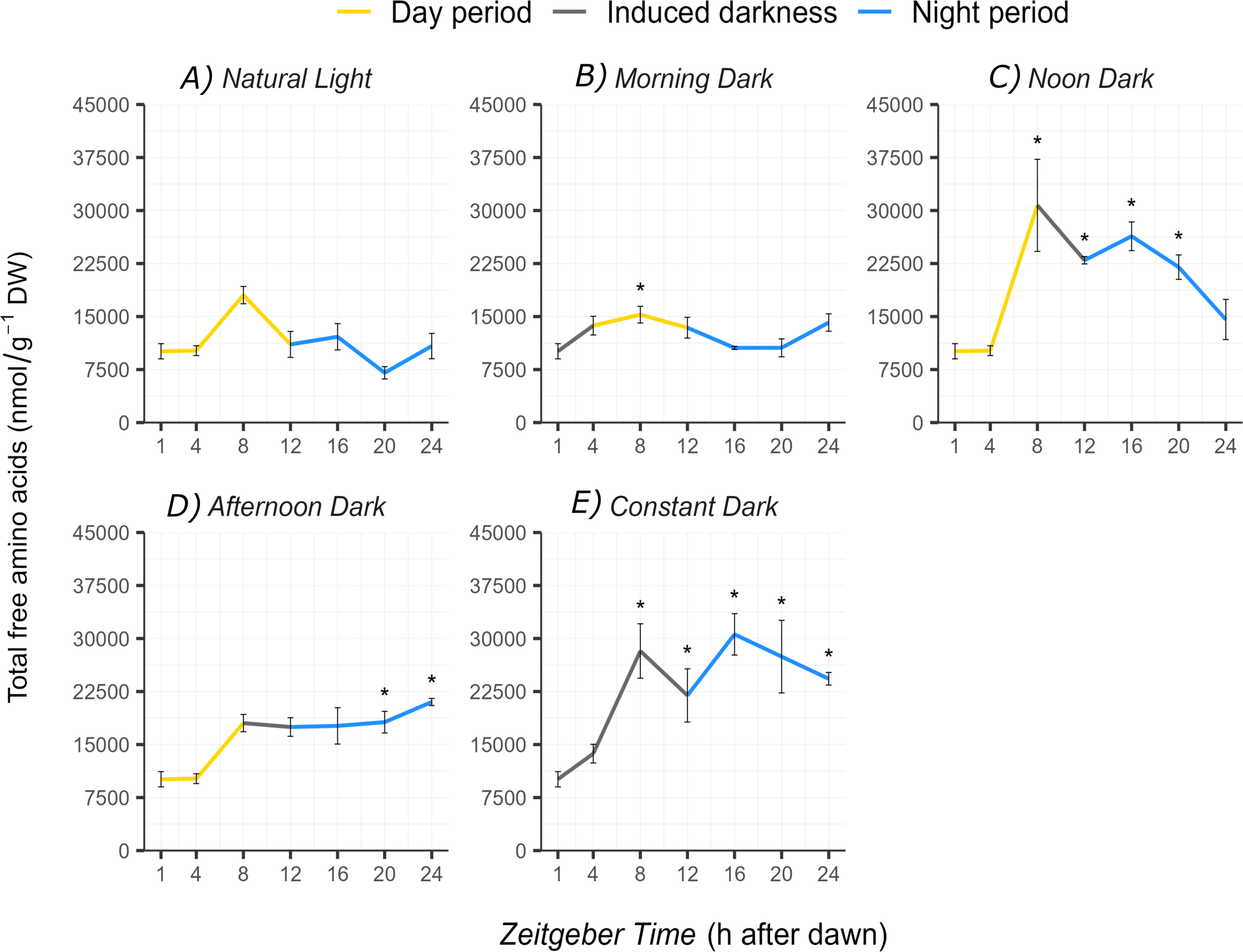
Diel variation in total free amino acid concentration in sugarcane leaf +1 under different periods of induced darkness. Total free amino acids were quantified at seven *Zeitgeber times* (ZT1, ZT4, ZT8, ZT12, ZT16, ZT20, and ZT24). ZT indicates the number of hours elapsed since dawn. Treatments comprised (A) Natural Light, (B) Morning Dark from ZT0 to ZT4, (C) Noon Dark from ZT4 to ZT8, (D) Afternoon Dark from ZT8 to ZT11, and (E) Constant Dark from ZT0 to ZT12. Yellow, grey, and blue line segments represent samples collected during the natural light period, induced darkness, and natural night, respectively. Concentrations are expressed as nmol mg⁻¹ dry weight. Values are means ± SE of five biological replicates. Asterisks indicate significant differences between each induced darkness treatment and Natural Light at the same ZT according to the Wilcoxon test at P < 0.05.

Noon Dark and Constant Dark showed the largest increases in total free amino acid concentrations, with values ranging from 17,481 to 30,590 nmol mg⁻¹ DW (Fig. 3). These changes became evident at ZT8 and persisted for much of the subsequent night. Most amino acid signals remained above those observed under Natural Light, although treatment-specific exceptions were observed for Asp, Gly, and Cit (Figs. S9 to S11). In Noon Dark, increases were particularly pronounced for branched-chain and aromatic amino acids, as well as Lys, Asn, Arg, and the combined Gln+His signal. Most of these amino acids peaked during the night and then declined toward ZT24. Leu, Ile, and Val showed particularly marked increases, indicating a strong reorganization of branched-chain amino acid pools following darkness imposed around midday.

### Diel circadian clock components maintain rhythmic expression under short-term dark induction

To assess whether darkness imposed at different times of day influences circadian regulation in sugarcane leaves, we examined diel expression of the core clock transcripts *ScLHY*, *ScTOC1*, *ScPRR37,* and *ScPRR73* across treatments (Fig. 4). Under Natural Light, *ScLHY* decreased from ZT1 to ZT8 and then rose again, reaching a secondary peak near ZT24, indicative of a morning-phased rhythm extending into the next dawn. A similar pattern was observed in Afternoon Dark. In Morning Dark, however, *ScLHY* recovery was delayed, with transcript levels remaining low until after ZT16. In Noon Dark, *ScLHY* expression decreased until ZT12 and then increased sharply by ZT24, whereas in Constant Dark it steadily declined up to ZT16, showing only partial recovery by the cycle’s end.

**Fig. 4.**
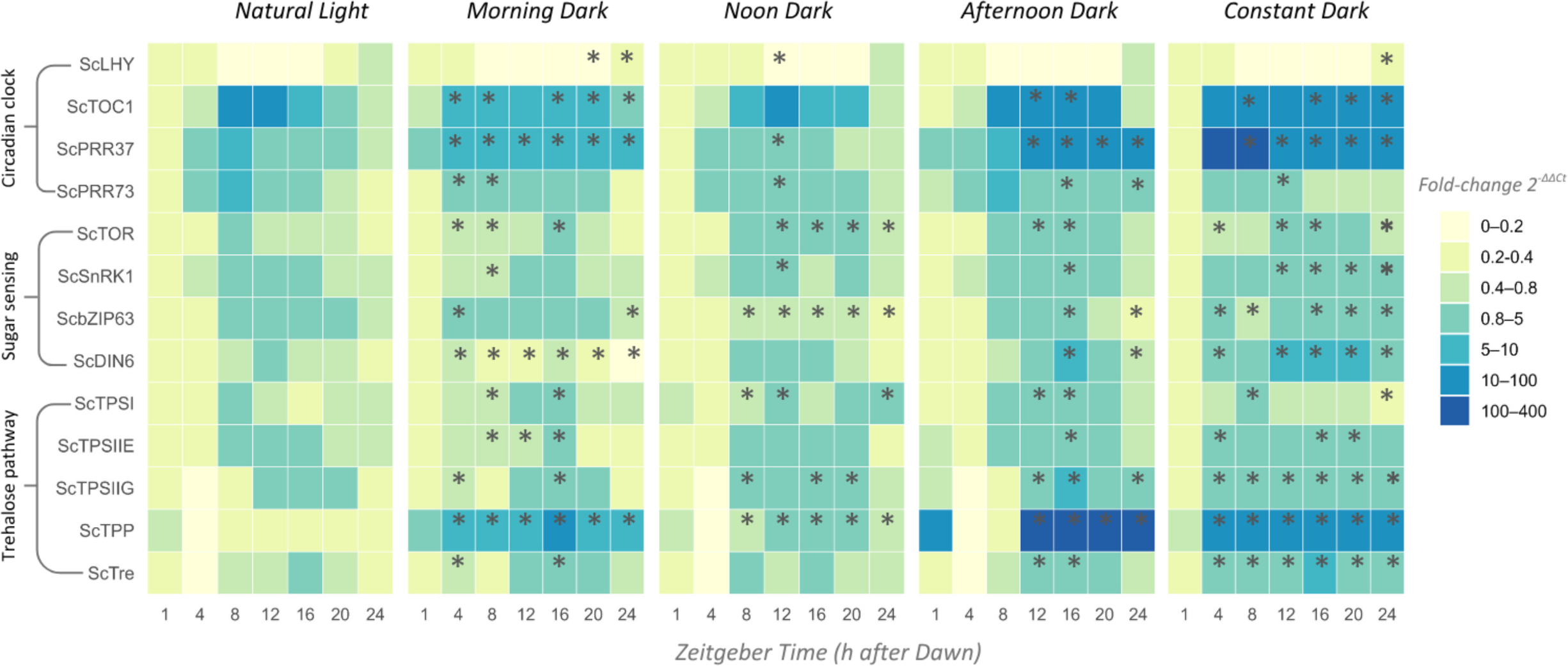
Diel expression profiles of clock associated, sugar sensing, low energy response, and trehalose pathway transcripts in sugarcane leaf +1 under different periods of induced darkness. Transcript abundance was analyzed for the clock associated genes *ScLHY*, *ScTOC1*, *ScPRR37*, and *ScPRR73*; the sugar sensing and low energy associated genes *ScTOR*, *ScSnRK1α*, *ScbZIP63*, and *ScDIN6*; and the trehalose pathway genes *ScTPSI*, *ScTPSIIE*, *ScTPSIIG*, *ScTPP*, and *ScTRE*. Samples were collected at seven *Zeitgeber times*, corresponding to ZT1, ZT4, ZT8, ZT12, ZT16, ZT20, and ZT24. ZT indicates the number of hours elapsed since dawn. Treatments comprised Natural Light, Morning Dark from ZT0 to ZT4, Noon Dark from ZT4 to ZT8, Afternoon Dark from ZT8 to ZT12, and Constant Dark from ZT0 to ZT12. Relative transcript abundance was calculated using the 2^−ΔΔCt^ method, with ZT1 as the calibrator, and normalized to the geometric mean of the reference genes *ScGAPDH*, *ScACT*, and *ScPGR*. Cell colors represent fold change according to the scale shown on the right. Yellow, grey, and blue bands above the heatmaps indicate the natural light period, induced darkness, and natural night, respectively. Values used to generate the heatmap are means of three biological replicates. Asterisks indicate significant differences between each induced darkness treatment and Natural Light at the same ZT according to the Wilcoxon test at P < 0.05.

*ScTOC1* exhibited the expected evening oscillation under Natural Light, with transcript levels rising after ZT4, peaking around ZT12, and declining during the night. In Morning Dark, *ScTOC1* was induced earlier, with higher levels already present at ZT4 and ZT8. Although expression declined after ZT12, it remained above control levels for the rest of the cycle. A similar advance in *ScTOC1* accumulation was observed in Afternoon Dark, whereas Constant Dark caused an early increase followed by stabilization at levels similar to those under Natural Light (Fig. S11).

The expression profile of *ScPRR37* varied substantially in amplitude across treatments, indicating activation of metabolic compensation mechanisms (Kugan *et al*., 2021). Under Natural Light, its highest transcript abundance was observed at ZT8 (Fig. S12). In Morning Dark and Constant Dark, expression increased significantly from ZT4 onward and remained elevated until ZT16 and ZT12, respectively. In contrast, Noon Dark and Afternoon Dark showed a delayed pattern, with the highest expression at ZT12. *ScPRR73* showed a more similar diel profile across treatments and reached its highest expression at ZT8 under Natural Light. However, the ZT8 increase was lower in all darkness treatments except Afternoon Dark.

### Sugar-sensing and energy-stress signaling transcripts respond dynamically to transient darkness

To investigate how transient darkness affects transcripts associated with sugar and energy signaling, we analyzed diel expression of *ScTOR*, *ScSnRK1*, *ScbZIP63*, and *ScDIN6* (Fig. 4). We also evaluated expression of five transcripts related to trehalose metabolism (*ScTPSI*, *ScTPSIIE*, *ScTPSIIG*, *ScTPP*, and *ScTRE*).

ScTOR transcript abundance increased after the onset of darkness across all induced-darkness treatments and peaked at approximately ZT16 (Fig. S13A). *ScSnRK1* expression also increased after dark exposure, although this response was not observed in Morning Dark (Fig. S13B). The temporal profiles of *ScbZIP63* and *ScDIN6* varied across treatments. The increase in *ScbZIP63* expression was less pronounced in Noon Dark, whereas *ScDIN6* showed a weaker response in Morning Dark (Fig. S13C and D). These patterns indicate that targets associated with low-energy signaling responded differently to darkness depending on the timing of the treatment.

Transcripts related to trehalose metabolism also exhibited treatment-dependent expression profiles. ScTPSI transcript abundance increased during darkness in the covered leaves, along with increased *ScTRE* expression. The two class II isoforms, *ScTPSIIE* and *ScTPSIIG*, showed broadly similar diel profiles and increased expression following dark exposure. The increase in ScTPSIIG was particularly pronounced in the Afternoon Dark. ScTPP expression also increased after the onset of induced darkness and during the natural night (Fig. S14).

### Free amino acids and trehalose pathway targets contribute to diel sample discrimination

To identify the variables that contributed most to diel variation within each treatment, we integrated physiological, metabolic, and transcript expression data using PLS-DA (Fig. 5). We built separate models for each treatment, with sampling time as the response class. We then used VIP scores to estimate each variable’s contribution to sample discrimination. The selected models achieved classification accuracies above 0.70, R² values above 0.80, and Q² values above 0.60 in fivefold cross-validation (Fig. S5). Permutation tests also indicated that the observed discrimination among sampling times exceeded chance expectations (P<0.001; Fig. S6).

**Fig. 5.**
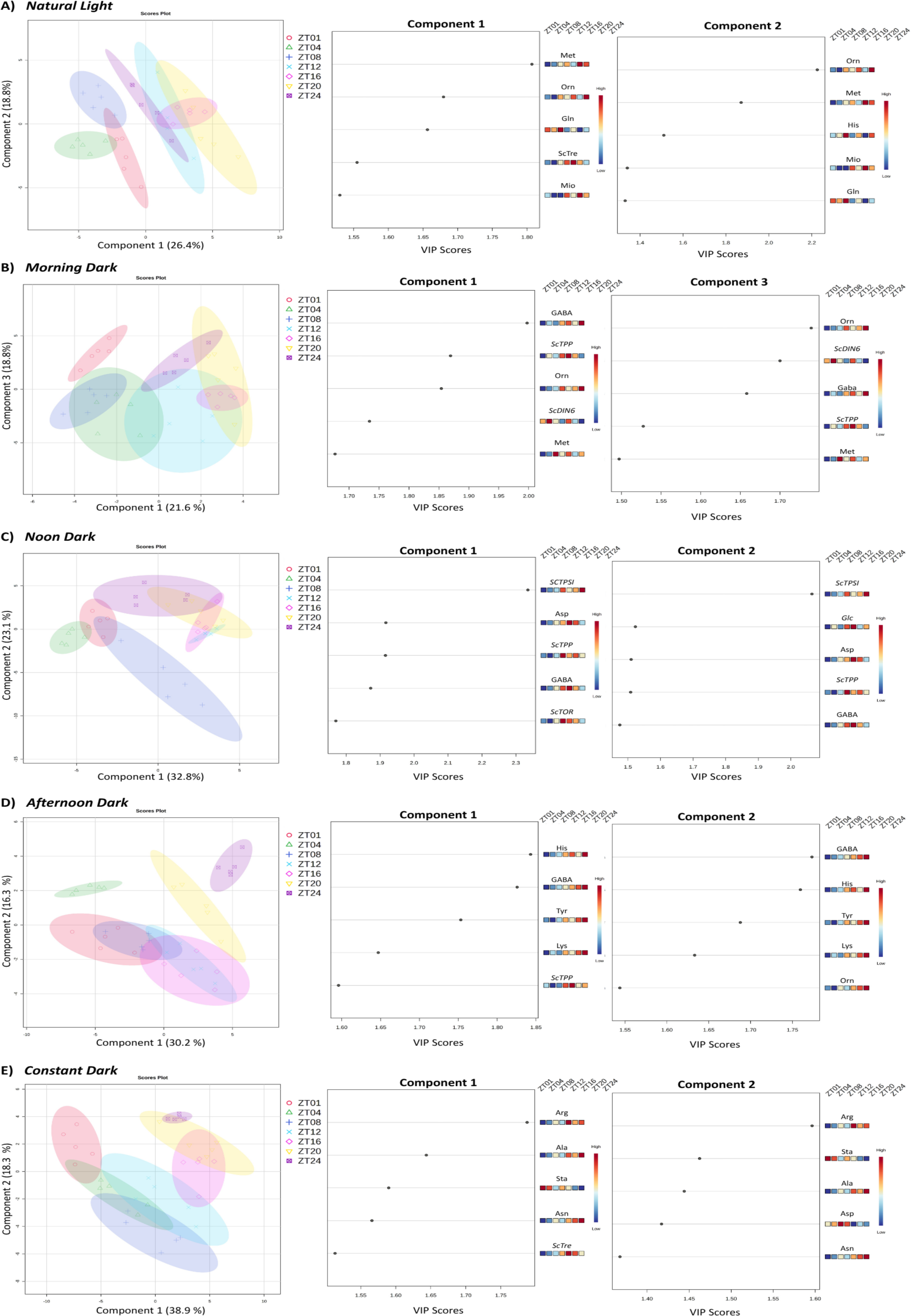
Partial least squares discriminant analysis (PLS-DA) of integrated physiological, metabolic, and transcriptional profiles in sugarcane leaf +1 throughout the diel cycle. Separate PLS-DA models were constructed for (A) Natural Light, (B) Morning Dark, (C) Noon Dark, (D) Afternoon Dark, and (E) Constant Dark, using *Zeitgeber time* as the response class. Each model included leaf gas exchange parameters, nonstructural carbohydrates, free amino acids, and relative transcript abundance of clock associated, sugar sensing, low energy response, and trehalose pathway genes. Score plots in the left panels show the distribution of biological samples collected at ZT1, ZT4, ZT8, ZT12, ZT16, ZT20, and ZT24 along the selected latent components. Shaded ellipses represent the distribution of samples from each sampling time. The percentage shown on each axis indicates the variation represented by the corresponding component. The center and right panels show Variable Importance in Projection (VIP) scores for the five variables contributing most strongly to each selected component. Adjacent heatmaps indicate the standardized relative abundance of each variable across sampling times, with blue and red representing lower and higher values, respectively.

Across treatments, samples collected during the daytime were generally separated from those collected during the natural night. Under Natural Light, the first two components accounted for 45.2% of the model variation. Component 1 accounted for 26.4% and primarily separated samples collected between ZT1 and ZT8 from those collected between ZT12 and ZT20. Met, Orn, and the combined Gln+His signal had among the highest VIP scores. Morning Dark showed a similar temporal separation. Met had the highest VIP score (1.85), while Orn, *ScTPP*, and the combined Tyr+GABA signal also contributed to sample discrimination. In Noon Dark, the first two components accounted for 55.9% of the model variation, with contributions of 32.8% and 23.1%, respectively. *ScTPSI* and *ScTPP expression*, along with Asp concentration, showed high VIP scores. The increase in *ScTPSI* expression during darkness coincided with changes in Asp concentration across the diel cycle (Fig. S9).

Under Afternoon Dark, component 1 accounted for 30.2% of the model variation. The combined Gln+His and Tyr+GABA signals were among the primary contributors to temporal discrimination. These amino acid signals peaked near ZT24 (Fig. S10). In Constant Dark, the first two components accounted for 57.2% of the model variation. Arg and Ala showed high VIP scores and generally increased across the cycle, whereas starch followed an opposing temporal pattern and progressively declined (Figs. 2 and 5).

### Correlation-based networks reveal treatment associated changes in physiological and metabolic covariation

To investigate associations among physiological, metabolic, and transcript expression variables, we constructed a correlation network for each treatment using Spearman’s rank correlation coefficients (Fig. 6). We retained edges when the absolute correlation coefficient exceeded 0.8. The resulting networks contained clusters composed predominantly of related variable groups, including amino acids, gas exchange parameters, and transcript expression profiles. Although this general organization was observed across treatments, differential network analysis detected changes in network topology relative to Natural Light (Table S2).

**Fig. 6.**
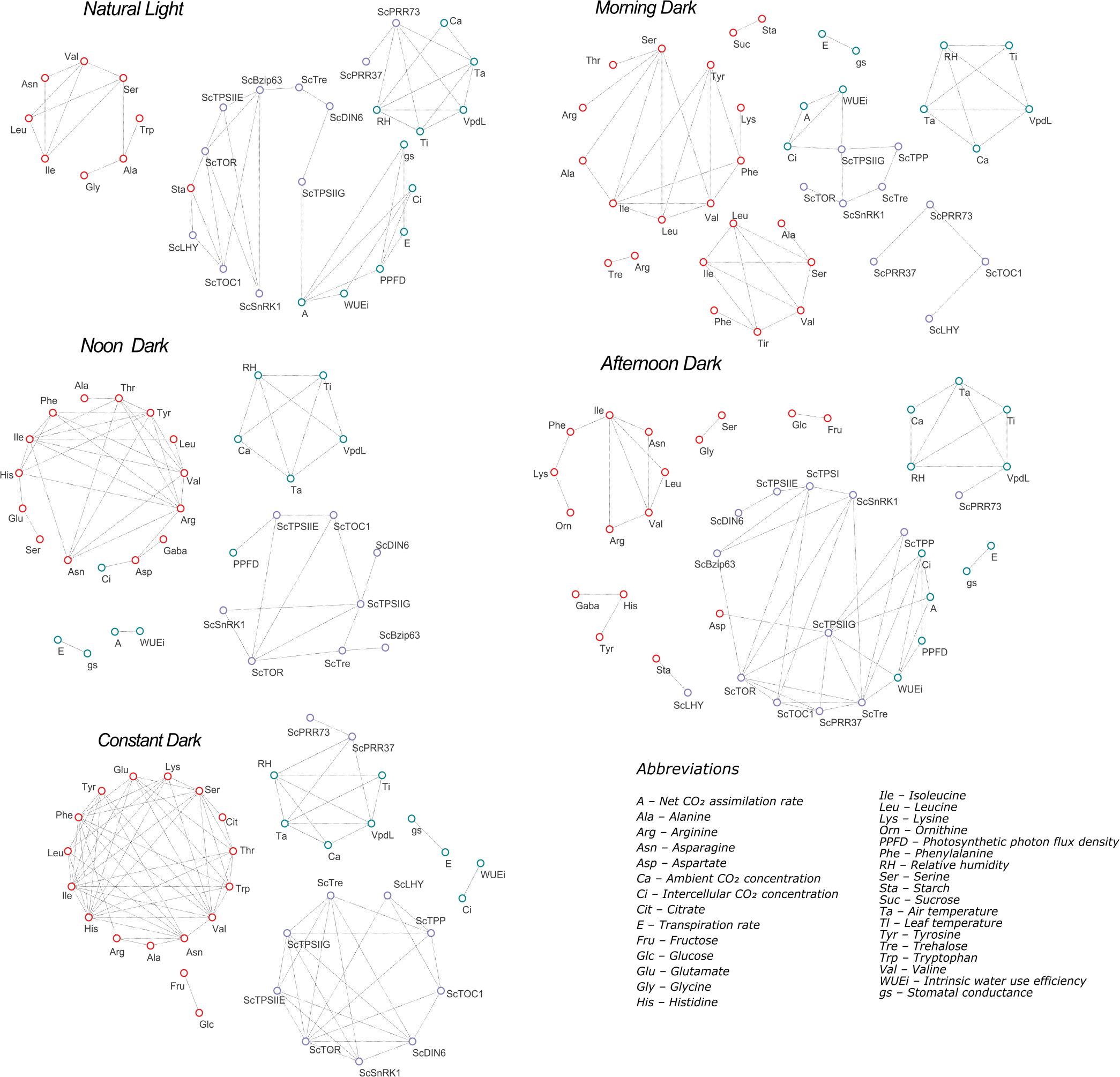
Co-variation networks integrating physiological, metabolic, environmental, and transcriptional variables in sugarcane leaf +1 throughout the diel cycle. Separate networks were constructed for (A) Natural Light, (B) Morning Dark, (C) Noon Dark, (D) Afternoon Dark, and (E) Constant Dark by combining data collected at ZT1, ZT4, ZT8, ZT12, ZT16, ZT20, and ZT24. Green, red, and purple nodes represent gas exchange and environmental variables, metabolites, and transcript abundance, respectively. Edges indicate associations retained according to Spearman’s rank correlation coefficient at ∣ρ∣>0.8. Only variables connected by at least one retained association are shown. Node positions were determined separately for each treatment to display their corresponding network organization. Networks were constructed using BioNetStat (Jardim et al., 2019) and visualized in Cytoscape version 3.8.0 (Shannon et al., 2003). Abbreviations are defined in the lower right panel.

Betweenness centrality differed significantly across the complete networks and the transcript and sugar-sensing subsets under all darkness treatments. Degree centrality detected fewer differences, most of which occurred in transcript-related subsets. Changes in closeness and eigenvector centrality were more frequent in metabolite-related subsets (Table S2). Together, these results indicate that associations among measured variables varied with the timing and duration of induced darkness.

Under Natural Light, the network contained three main clusters. One cluster was dominated by amino acids, another by environmental variables and clock-associated transcripts, and a third by sugar-sensing transcripts and gas-exchange parameters. Within this network, *ScTPSIIG* was negatively correlated with net CO₂ assimilation and associated with variables from different functional groups (Fig. S15). Starch was negatively correlated with *ScLHY* and positively correlated with *ScTOC1*. The nodes with the highest degree centrality were *ScTOR*, *ScPRR37*, *ScbZIP63*, and net CO₂ assimilation, each with five connections.

Associations between transcript expression and physiological or metabolic variables shifted after induced darkness. Clock-associated transcripts retained several correlations with one another, but their associations with metabolites differed across treatments. This was particularly evident in correlations involving starch. Among the clock-associated transcripts, *ScTOC1* showed the largest changes in connectivity relative to Natural Light (Table S3). *ScTOR* and *ScTPSIIG* connectivity also varied across treatments, and both transcripts remained among the most highly connected transcript nodes. These patterns identify *ScTOC1*, *ScTOR*, and *ScTPSIIG* as candidate nodes associated with the network response to transient darkness.

In Morning Dark, *ScTPSIIG* was negatively correlated with intrinsic water use efficiency and positively correlated with intercellular CO₂ concentration. The association between *ScTPSIIG* and stomatal conductance observed under Natural Light was no longer present. Sucrose and starch were positively correlated, while the clock-associated genes formed a cluster with positive correlations between the two *ScPRR* transcripts and a negative correlation between *ScTOC1* and *ScLHY*. The nodes with the highest degree included *ScTPSIIG*, Ser, the combined Tyr+GABA signal, and Leu.

Afternoon Dark also altered ScTPSIIG associations with transcript expression and gas exchange variables. In this treatment, *ScTOC1* and *ScPRR37* were positively correlated, whereas *ScTRE*, *ScTOR*, and *ScTPSIIG* showed the highest degree centrality, each with seven connections. Under Noon Dark and Constant Dark, we observed fewer associations between transcript expression and gas exchange variables. In contrast, amino acids formed a more densely connected cluster that included Asn, Asp, Glu, Ile, and the combined Gln+His signal (Table S3).

## Discussion

### The timing of transient darkness determines the magnitude and nature of the physiological perturbation

Under field conditions, sugarcane leaves exhibit a clear diel pattern of CO₂ assimilation that tracks daily changes in irradiance, temperature, and atmospheric demand (De Souza et al., 2018). Variations in light intensity and spectral composition from cloud cover, canopy structure, and local microclimatic conditions also influence photosynthetic performance (Chiang et al., 2019; Dantas et al., 2021). Our results show that suppressing photosynthetic carbon assimilation at different times of day produced distinct physiological and metabolic responses. These differences indicate that the consequences of transient darkness depend on the leaf’s physiological and metabolic state when light is interrupted.

The effects of darkness were particularly evident when the treatment overlapped with the period of highest photosynthetic activity, from ZT4 to ZT8. Under Natural Light, this interval encompassed the daily peak in CO₂ assimilation and preceded the strongest accumulation of sucrose and starch (De Souza et al., 2018; de Oliveira et al., 2025). Darkness imposed around midday prevented carbon assimilation during this period and was followed by marked changes in carbohydrate and amino acid profiles. The Noon Dark network also differed from that observed under Natural Light, particularly in associations involving gas exchange variables and metabolites (Fig. 6 and Table S2). These findings suggest that interrupting photosynthesis near its daily maximum has consequences that persist after light exposure is restored. However, comparisons with Constant Dark should be interpreted cautiously because this treatment differed in both the timing and duration of darkness.

Darkness imposed at dawn produced a distinct response. Net CO₂ assimilation remained lower than under Natural Light after reillumination, indicating incomplete photosynthetic recovery during the remaining photoperiod (Fig. 1). This response coincided with changes in the association between stomatal conductance and PPFD (Fig. S15), which may reflect altered stomatal responsiveness following the absence of the morning light signal. Stomatal opening generally responds more slowly than photosynthetic biochemistry to changes in irradiance (McAusland et al., 2016). Its induction at dawn also depends on light perception, particularly blue light signaling (Inoue and Kinoshita, 2017). When illumination was restored later in the morning, the higher leaf-to-air vapor pressure deficit may have further limited stomatal conductance and delayed the recovery of CO₂ assimilation (De Souza et al., 2018).

Despite lower carbon assimilation, Morning Dark leaves accumulated relatively high concentrations of sucrose and starch after reillumination (Fig. 2). This apparent discrepancy suggests that carbohydrate concentrations were influenced not only by current carbon assimilation but also by differences in carbon utilization and export. Reduced sucrose export or lower carbon demand may have contributed to sucrose retention in the leaf, although neither process was measured directly. Changes in carbohydrate export under contrasting light conditions have also been reported in grapevine, indicating that carbon availability can modify the balance between leaf retention and export to sink tissues (Dayer et al., 2021). Starch accumulated after reillumination and then declined during the night, while soluble sugar concentrations remained comparatively high. This pattern is consistent with transient starch reserves contributing to nocturnal carbon availability.

The relationship between starch and clock-associated transcript expression also varied across treatments. Under Natural Light, starch was negatively correlated with *ScLHY* and positively correlated with *ScTOC1* (Fig. S15), consistent with the established association between diel carbon metabolism and the temporal regulation of starch turnover (Hotta et al., 2013; Dantas et al., 2020). These correlations shifted after induced darkness, indicating that the temporal relationship between starch concentration and clock-associated transcription was sensitive to light interruption.

Afternoon Dark further showed that a late interruption of photosynthesis reduced carbohydrate availability during the subsequent night. Although starch reached concentrations like those under Natural Light before nightfall, both starch and sucrose declined more sharply toward ZT24, while free amino acids accumulated. These patterns suggest that the timing of transient darkness influenced the amount and composition of carbon reserves available during the night. Taken together, the contrasting responses to Morning, Noon, and Afternoon Dark indicate that sugarcane leaves do not respond uniformly to short interruptions in photosynthetic carbon supply. Instead, the metabolic consequences depend on when the interruption occurs and on the carbohydrate pools available during the subsequent recovery and night periods.

### Residual sucrose availability distinguishes buffered metabolic adjustment from carbon-starvation associated responses

Transient reductions in photosynthetic carbon supply can disrupt the balance between carbon and nitrogen metabolism and promote a shift from biosynthetic activity toward energy conservation (Araújo et al., 2010; Law et al., 2018). However, our results indicate that the response to induced darkness depended not only on interrupted photosynthesis but also on the carbohydrate pools available during the subsequent diel cycle. Rather than producing a uniform starvation response, the treatments generated distinct metabolic profiles that differed in sucrose availability, starch concentration, amino acid accumulation, and expression of low-energy-associated genes (Kerr et al., 1985; Fujiki et al., 2001).

This contrast was particularly evident between Noon Dark and Constant Dark. Both treatments maintained low starch concentrations and promoted free amino acid accumulation during the diel cycle (Figs. 2 and 3). Starch serves as an important carbon source when photosynthetic assimilation ceases and soluble sugars become limiting, supporting metabolism throughout the night (Ishizaki et al., 2005; Araújo et al., 2010). Despite similarly low starch pools, Noon Dark and Constant Dark differed markedly in sucrose dynamics. Sucrose recovered after reillumination and remained relatively stable in Noon Dark, whereas it progressively declined in Constant Dark. These contrasting profiles indicate that amino acid accumulation alone is insufficient to define the severity of carbon limitation. Instead, sucrose maintenance after reillumination may have moderated the response in Noon Dark, whereas its continued depletion under Constant Dark was associated with a stronger starvation-related profile.

Changes in individual amino acids further supported this distinction. Several treatments showed increased concentrations of Ala, Asp, Arg, Glu, and the combined Gln+His signal during or after induced darkness (Figs. S7 to S11). Amino acids can accumulate under carbon limitation through shifts in protein degradation, biosynthesis, catabolism, transport, and incorporation into proteins (Hildebrandt et al., 2015; Batista-Silva et al., 2019). Their accumulation may also provide alternative respiratory substrates when carbohydrate availability is limited (Ishizaki et al., 2005; Izumi et al., 2013). In particular, Arg and Gln have been associated with nitrogen storage and subsequent remobilization during recovery from stress (Batista-Silva et al., 2019; McLoughlin et al., 2020).

In Noon Dark, free amino acid concentrations increased at night even though sucrose had recovered after reillumination. This pattern may reflect a persistent metabolic effect of the preceding interruption in photosynthesis rather than an irreversible transition to carbon starvation. The accumulated amino acid pool could contribute to subsequent carbon and nitrogen remobilization, while maintaining sucrose could preserve a readily available carbon source. These complementary responses may have allowed the leaf to adjust its metabolism without adopting the stronger low-energy profile observed under Constant Dark.

The expression profiles of low-energy-associated transcripts also differed between the two treatments (Fig. 4). Under Constant Dark, the progressive decline in sucrose coincided with increased transcript abundance of *ScSnRK1*, *ScbZIP63*, and *ScDIN6*. Together with the accumulation of free amino acids, this pattern is consistent with enhanced SnRK1-associated low-energy signaling (Baena-González et al., 2007; Margalha et al., 2019). In Noon Dark, *ScbZIP63* and *ScDIN6* remained at or below the levels observed under Natural Light, despite the interruption of photosynthetic assimilation. These contrasting responses suggest that sucrose preservation was associated with a weaker transcriptional response to low-energy status.

The relationship between *ScDIN6* expression and amino acid profiles further supports this interpretation. SnRK1 can phosphorylate bZIP63, which regulates low-energy-responsive genes, including DIN6/ASN1 (Matiolli et al., 2011; Frank et al., 2018). In Constant Dark, *ScSnRK1* and *ScDIN6* expression were positively correlated, and *ScbZIP63* transcript abundance also increased (Fig. S15). Higher *ScDIN6* expression coincided with increased Asn and lower Asp concentrations, consistent with altered Asn metabolism via the ASN1 pathway (Lam et al., 2003; Law et al., 2018). In Noon Dark, Asp remained more abundant than Asn, and *ScDIN6* expression showed a weaker response (Fig. S9). This difference is consistent with the known repression of *DIN* genes by sucrose (Lam et al., 1994, 2003).

Taken together, these results support a model in which residual sucrose availability distinguishes a buffered metabolic adjustment from a stronger starvation-associated response. In Noon Dark, sucrose recovery was associated with a limited transcriptional response in *ScbZIP63* and *ScDIN6*, despite pronounced changes in amino acid pools. In Constant Dark, continued sucrose depletion coincided with broader amino acid accumulation and increased expression of transcripts associated with low-energy signaling. Residual sucrose therefore appears to reflect the metabolic state reached after transient or prolonged darkness.

### TPS class II transcripts emerge as candidate nodes in the response to transient darkness

The contrasting responses across treatments indicate that sugarcane leaves adjust carbon assimilation, carbohydrate storage, and metabolic demand in response to the timing of light availability. During the transition from day to night, plants must balance sucrose export with accumulating sufficient carbon reserves to sustain metabolism in darkness (Stitt et al., 2010). Coordinated changes in carbohydrate pools, amino acid concentrations, transcript abundance, and gas exchange suggest that the response to transient darkness involves several interconnected physiological and metabolic processes (Henry et al., 2014; Lunn et al., 2014; Figueroa et al., 2016). In this context, components of the trehalose pathway emerged as candidates for linking carbon status to broader adjustments in leaf metabolism.

Multivariate analyses consistently identified trehalose pathway transcripts and amino acids as the variables that contributed most strongly to temporal sample discrimination (Fig. 5). Free amino acids and transcripts such as *ScTPSI* and *ScTPP* frequently showed high VIP scores, though their relative contributions varied across treatments. Under Natural Light, Met, Orn, and the combined Gln+His signal were among the main discriminating variables. Under Morning Dark, Met, Orn, *ScTPP*, and the combined Tyr+GABA signal contributed strongly to temporal separation. Under Noon Dark, *ScTPSI*, *ScTPP*, and Asp were prominent, whereas Arg, Ala, and starch contributed strongly under Constant Dark. These results indicate that amino acid pools and trehalose pathway transcription were important components of diel metabolic variation under both natural and induced darkness.

The correlation networks provided complementary information, showing that associations among physiological, metabolic, and transcriptional variables differed across treatments (Fig. 6 and Tables S2 and S3). Several sugar-sensing transcripts changed their connectivity and topological position, suggesting that transient darkness affected patterns of covariation beyond transcript abundance. Betweenness centrality showed consistent treatment-associated changes in the complete networks and in gene-related subsets. Degree centrality detected additional changes involving sugar-sensing transcripts, whereas closeness and eigenvector centralities were more responsive in metabolite-related subsets. These topological differences do not demonstrate regulatory interactions, but they identify candidate components associated with the integrated response to transient darkness.

The phylogenetic positions of the two sugarcane class II TPS isoforms provide an important basis for interpreting their potentially distinct roles. The plant TPS family comprises class I and class II proteins, corresponding to clades B and A, respectively. Class II proteins are further distributed across several subclades (de Oliveira et al., 2022). *ScTPSIIE* belongs to subclade A3 and is phylogenetically related to *A. thaliana TPS7*, whereas *ScTPSIIG* belongs to subclade A2 and is related to *AtTPS6*. This distinction suggests that the two sugarcane isoforms represent distinct regulatory lineages rather than interchangeable components of the trehalose pathway.

The relationship between *ScTPSIIE* and *AtTPS7* is particularly relevant to the changes in amino acid metabolism and water-related traits observed in our study. Loss of *AtTPS7* affects amino acid accumulation and energy-related metabolites during stress recovery. It also alters responses associated with ABA, stomatal regulation, photosynthesis, and water use efficiency (Bonilla-Córdoba et al., 2026). Under Constant Dark, increased *ScTPSIIE* expression coincided with pronounced accumulation of free amino acids. Expression also changed under Morning Dark, when stomatal conductance and carbon assimilation showed incomplete recovery after reillumination. These similarities are consistent with a possible role for the A3 lineage in coordinating metabolic and water-related responses.

*ScTPSIIG*, which belongs to the *AtTPS6*-related A2 subclade, showed the clearest treatment-associated changes in network position. Under Natural Light, *ScTPSIIG* connected sugar-sensing transcripts with physiological variables and was negatively correlated with net CO₂ assimilation. Under Morning Dark, it was negatively correlated with intrinsic water use efficiency and positively correlated with intercellular CO₂ concentration, and it no longer associated with stomatal conductance. Under Afternoon Dark, *ScTPSIIG* again connected transcriptional and gas exchange variables and shared the highest degree centrality with *ScTRE* and *ScTOR* (Figs. 6 and S14). These patterns indicate that *ScTPSIIG* remained associated with carbon assimilation and water-related traits, although its connectivity varied with darkness timing.

Placing *ScTPSIIG* in the A2 subclade, together with *AtTPS6,* provides additional context for these associations. *AtTPS6* has been implicated in regulating cell morphology and plant architecture, and experimental evidence suggests it may retain unusual biochemical properties compared with other class II TPS proteins (Chary et al., 2008). The phylogenetic relationship with *AtTPS6* therefore supports a broader role for *ScTPSIIG* in sugar-related and physiological responses. However, this relationship does not demonstrate that the sugarcane protein shares the biochemical activity or cellular functions reported for *AtTPS6*. Experimental determination of ScTPSIIG’s catalytic properties and physiological function is needed.

The relative contributions of trehalose pathway transcripts and amino acids also differed across treatments. Under Morning and Afternoon Dark, *ScTPSIIG* and other sugar-sensing transcripts retained several associations with gas exchange variables. Under Noon Dark and Constant Dark, amino acids formed a more densely connected group, particularly through associations involving Ala, Asp, Arg, Asn, Glu, Ile, and the combined Gln+His signal. This pattern mirrored the high VIP scores of several amino acids in the corresponding PLS-DA models and their accumulation under treatments that strongly restricted photosynthetic assimilation. These results indicate that coordinated variation in amino acid pools became more prominent as carbohydrate availability declined.

The distinct responses of *ScTPSIIE* and *ScTPSIIG* indicate that class II TPS genes should not be treated as a functionally uniform group. *ScTPSIIE* was more closely associated with amino acid accumulation under Constant Dark, whereas *ScTPSIIG* showed broader associations with gas exchange, water-use traits, sugar-sensing genes, and other components of the trehalose pathway. These differences align with their placement in the A3 and A2 subclades, respectively, and suggest that the two isoforms may contribute to different aspects of the response to transient darkness.

Class II TPS proteins may represent candidate molecular links between these transcriptional and metabolic responses. Unlike class I TPS enzymes, most class II proteins are considered to have predominantly regulatory functions (Figueroa and Lunn, 2016). Recent evidence indicates that class II TPS proteins can interact with SnRK1 and may connect sugar availability to low-energy signaling (Van Leene et al., 2022; Reis-Barata et al., 2025, Preprint). Interactions with catalytic components of the trehalose pathway have also been proposed (Chary et al., 2008; Zang et al., 2011). Consistent with this framework, *ScTPSIIE* and *ScTPSIIG* were correlated with *ScTPSI*, *ScTPP*, and *ScTRE* across several treatments (Fig. S14). These associations support their selection as candidate components of the sucrose, Tre6P, and SnRK1 signaling framework.

Taken together, the results support a working model in which the timing of transient darkness influences the carbohydrate pools available during recovery and the subsequent night. Residual sucrose was associated with the intensity of the low-energy response, whereas amino acid pools and trehalose pathway transcripts contributed strongly to temporal discrimination and network organization. *ScTPSIIE* was mainly associated with amino acid accumulation and water-related responses. *ScTPSIIG* showed broader connections with carbon metabolism, gas exchange, sugar sensing, and other trehalose pathway transcripts. These convergent patterns identify both isoforms as candidates for functional investigation.

## Conclusion

Our findings show that sugarcane integrates light cues, energy status, and sugar-sensing circuits in a strongly time-of-day-dependent manner, revealing that diel metabolic coordination is more dynamic and more sensitive to carbon deprivation than previously recognized in C₄ species (Fig. 7). Transient darkness induced both stomatal and metabolic limitations that hindered photosynthetic reactivation and disrupted the natural relationship among carbohydrate export, starch turnover, and circadian coupling. Under these conditions, plants switched between sucrose-preserving strategies and more intense catabolic responses, as evidenced by restructuring of amino acid profiles and selective activation of SnRK1 and TOR modules. The balance between amino acid mobilization and metabolic regulation was closely associated with residual sucrose availability, identifying this sugar as a key indicator of the metabolic state reached under carbon limitation. We also observed significant reorganization of the trehalose pathway, in which class II TPS isoforms emerged as candidate links associated with carbon status, water balance, nitrogen metabolism, and Tre6P–SnRK1 signaling. Overall, our study improves understanding of metabolic resilience in a large-statured C₄ crop and highlights components of the trehalose pathway as promising targets for functional investigation of photosynthetic stability and carbon-use efficiency under varying light and energy conditions.

**Fig. 7.**
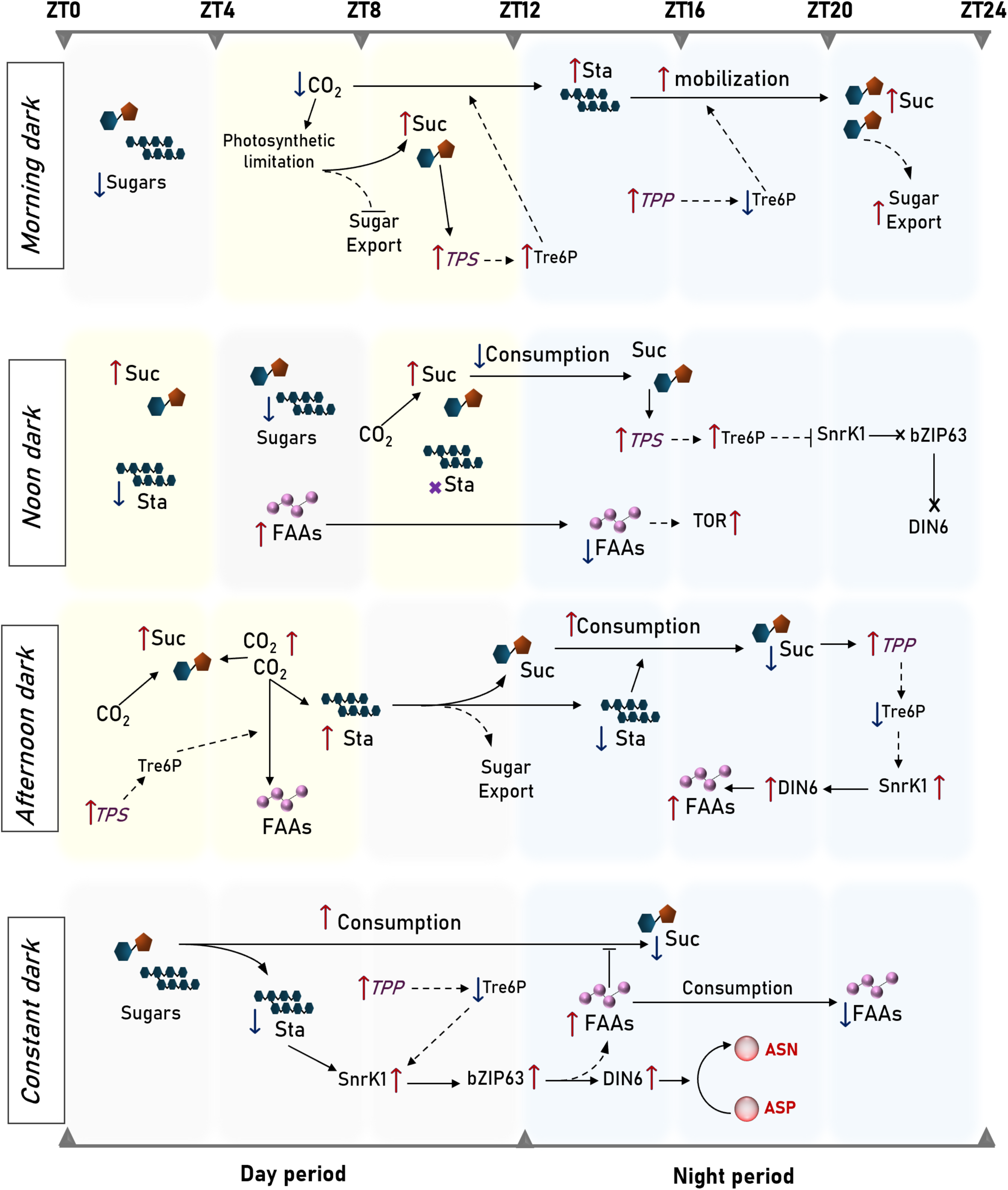
Conceptual model of time dependent physiological, metabolic, and transcriptional responses to induced darkness in sugarcane leaf +1. The model summarizes the main changes observed during a 24 h diel cycle, expressed as *Zeitgeber time* from ZT0 to ZT24. The natural light and night periods are represented by pale yellow and pale blue backgrounds, respectively. Rows correspond to Morning Dark from ZT0 to ZT4, Noon Dark from ZT4 to ZT8, Afternoon Dark from ZT8 to ZT12, and Constant Dark from ZT0 to ZT12. Changes in soluble sugars, sucrose, starch, free amino acids, CO₂ assimilation, and transcript abundance are shown along the diel cycle. Red and blue arrows indicate increases and decreases relative to Natural Light at the corresponding sampling time. Purple labels indicate changes in transcript abundance of genes associated with the trehalose pathway and low energy signalling. Solid arrows represent measured temporal changes or established metabolic relationships, while dashed arrows represent regulatory relationships proposed from the integrated interpretation of transcript expression, metabolite profiles, PLS-DA, and correlation networks. The model highlights the distinct responses associated with the timing of induced darkness. These include incomplete photosynthetic recovery after Morning Dark, preservation of sucrose despite low starch under Noon Dark, stronger nocturnal depletion of carbohydrate pools after Afternoon Dark, and the development of a starvation associated profile under Constant Dark. Changes in trehalose pathway transcripts and in the *SnRK1α*, *bZIP63*, and *DIN6* associated response are presented as proposed components of these metabolic adjustments. Abbreviations include ZT for *Zeitgeber time*, Suc for sucrose, Sta for starch, Tre6P for trehalose 6-phosphate, TPS for trehalose 6-phosphate synthase, TPP for trehalose 6-phosphate phosphatase, FAAs for free amino acids, SnRK1 for SNF1 related protein kinase 1, TOR for Target of Rapamycin, bZIP63 for basic leucine zipper 63, DIN6 for dark inducible 6, Asn for asparagine, and Asp for aspartate.

## Author contributions

**Gabriel Marques Leal:** Conceptualization, Formal Analysis, Investigation, Methodology, Validation Writing – Original Draft. **Hellen Oliveira de Oliveira:** Investigation, Methodology, Validation. **Felipe Farineli:** Data Curation, Methodology. **Arthur Vanni-Lopes:** Investigation, Methodology. **William Vinícius de Mello Mira:** Investigation, Methodology. **Amanda Ferreira Macedo:** Investigation, Methodology. **Grayce Hellen Romim:** Investigation. **Carlos Takeshi Hotta:** Methodology, Resources, Writing – Review & Editing. **Eny Iochevet Segal Floh:** Methodology. **Bruno Viana Navarro:** Conceptualization, Investigation, Methodology, Validation, Writing – Review & Editing. **Marcos Silveira Buckeridge:** Conceptualization, Project Administration, Supervision, Writing – Review & Editing

## Acknowledgements

We thank the University of São Paulo, the National Council for Scientific and Technological Development (CNPq), the São Paulo Research Foundation (FAPESP), and the Coordination for the Improvement of Higher Education Personnel (CAPES) for supporting this study.

