## Supplementary Figures for "Timing of transient darkness shapes carbon–nitrogen metabolism and sugar signaling in sugarcane"

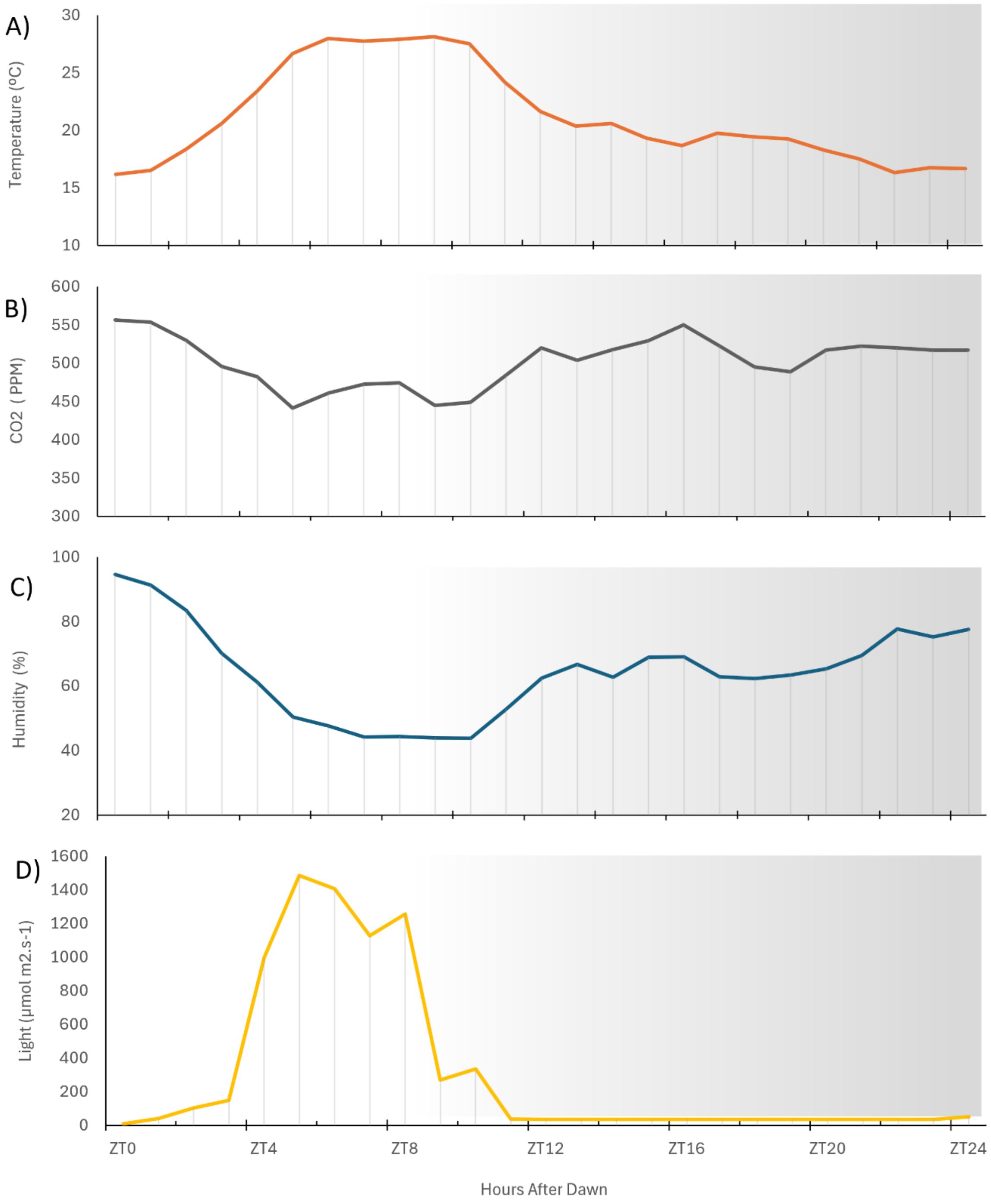


**Fig. S1. Environmental conditions in the experimental area during the diel cycle.** Variations in (A) air temperature, (B) ambient $CO_2$, (C) relative humidity and (D) photosynthetically active radiation recorded in the experimental area of the Laboratório de Fisiologia Ecológica de Plantas, (LAFIECO, Universidade de São Paulo, Brazil) from dawn on 23 May (ZT0) to dawn on 24 May 2024 (ZT24).

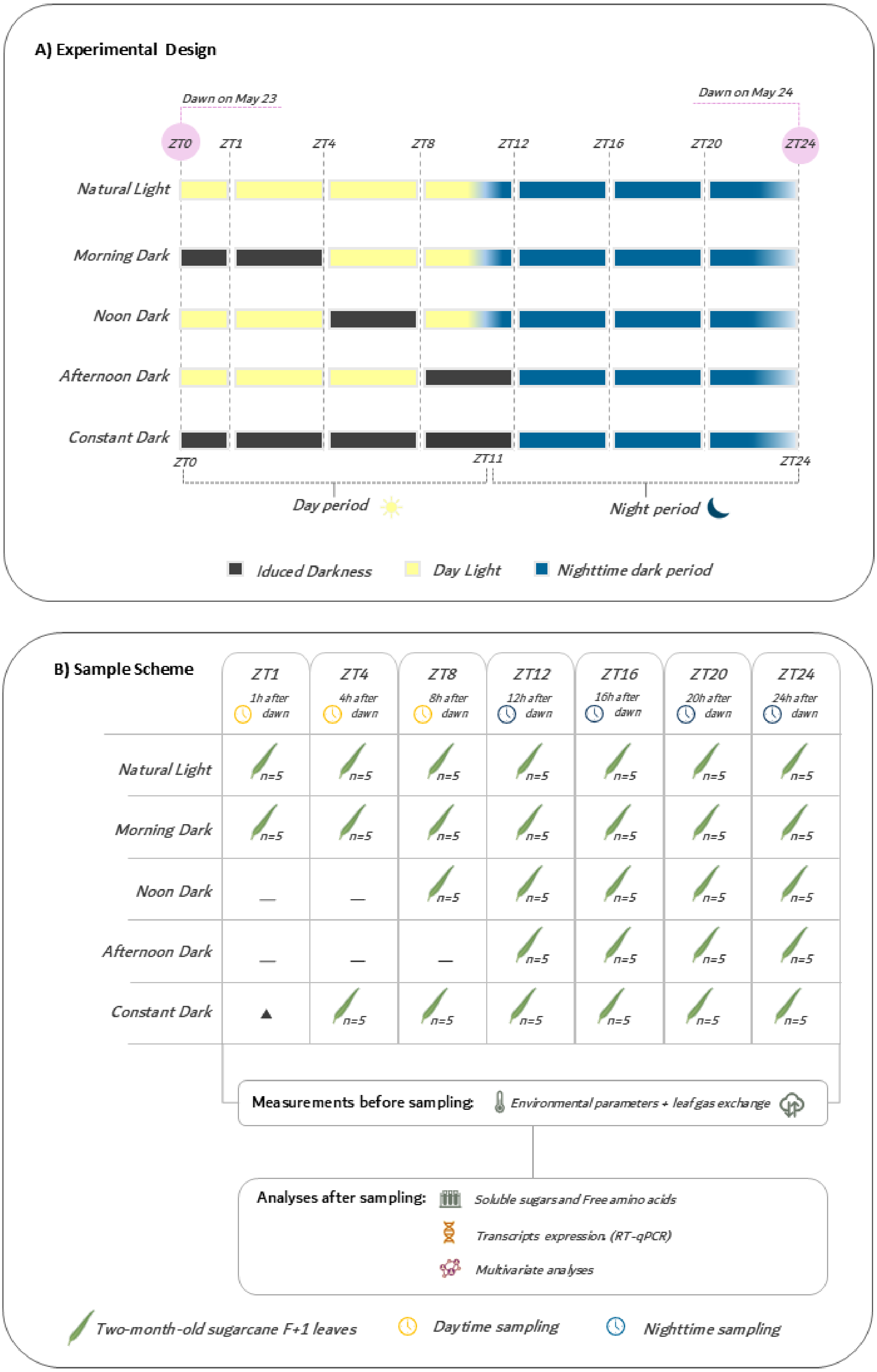


**Fig. 2. Experimental design and sampling scheme for diel characterization of physiological, metabolic, and transcriptional responses to induced T=transient darkness in sugarcane leaf +1.** ***(A)*** Experimental design showing the 24 h diel cycle expressed as *Zeitgeber time* (ZT), from ZT0 (dawn, May 23, 2024) to ZT24 (dawn, May 24, 2024). The natural light period extended from ZT0 to ZT11, followed by the nocturnal dark period from ZT11 to ZT24. Five darkness treatments were imposed on leaf +1: Natural Light, Morning Dark (ZT0–ZT4), Noon Dark (ZT4–ZT8), Afternoon Dark (ZT8–ZT12), and Constant Dark (ZT0–ZT12). Black, yellow, and blue bars indicate induced darkness, daylight, and

nighttime, respectively. ***(B)*** Sampling scheme and downstream analyses. Two-month-old sugarcane F+1 leaves were collected at ZT1, ZT4, ZT8, ZT12, ZT16, ZT20, and ZT24, with five biological replicates per group. The same set of five plants was used to represent treatments sharing identical light conditions at a given time point. (—) indicates use of the Natural Light control set; (▲) indicates use of the Morning Dark set. Leaf gas exchange and environmental parameters were monitored throughout the diel cycle prior to sampling. Collected leaves were analyzed for soluble sugars, free amino acids, and transcript abundance by RT-qPCR. Resulting datasets were further evaluated using multivariate, correlation, and network analyses.

*A) Natural Light*

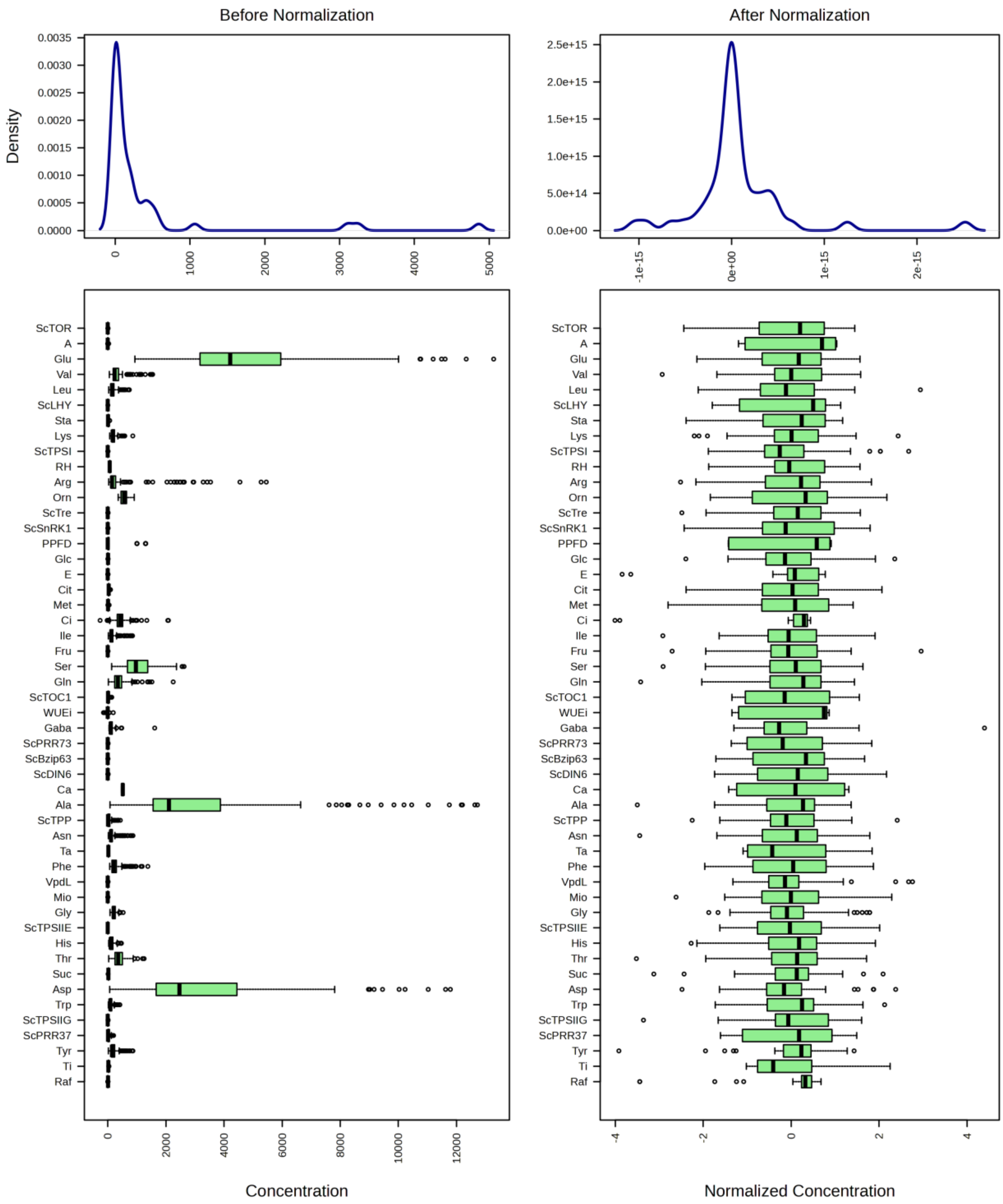

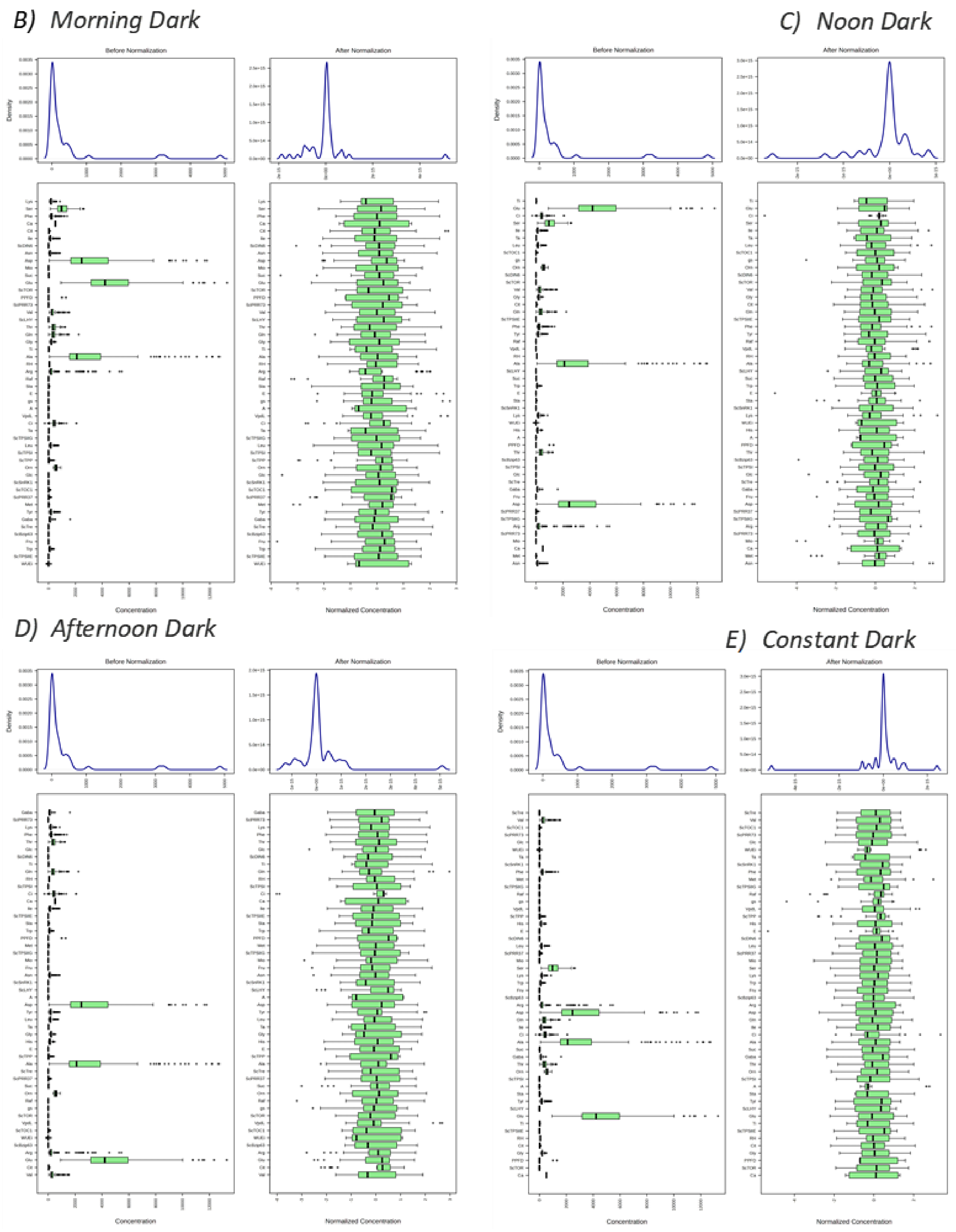


**Fig. S3. Distribution and normalization of the variables included in the PLS-DA analysis.** Density plots (top) and boxplots (bottom) illustrate the distribution of the original variable values (left) and the normalized data (right), obtained after $\log_{10}$ transformation, mean centering, and scaling to unit variance in MetaboAnalyst 5.0. The normalization procedure was applied to all 51 variables across samples from the five darkness treatments: (A) Natural Light, (B) Morning Dark, (C) Noon Dark, (D) Afternoon Dark, and (E) Constant Dark.

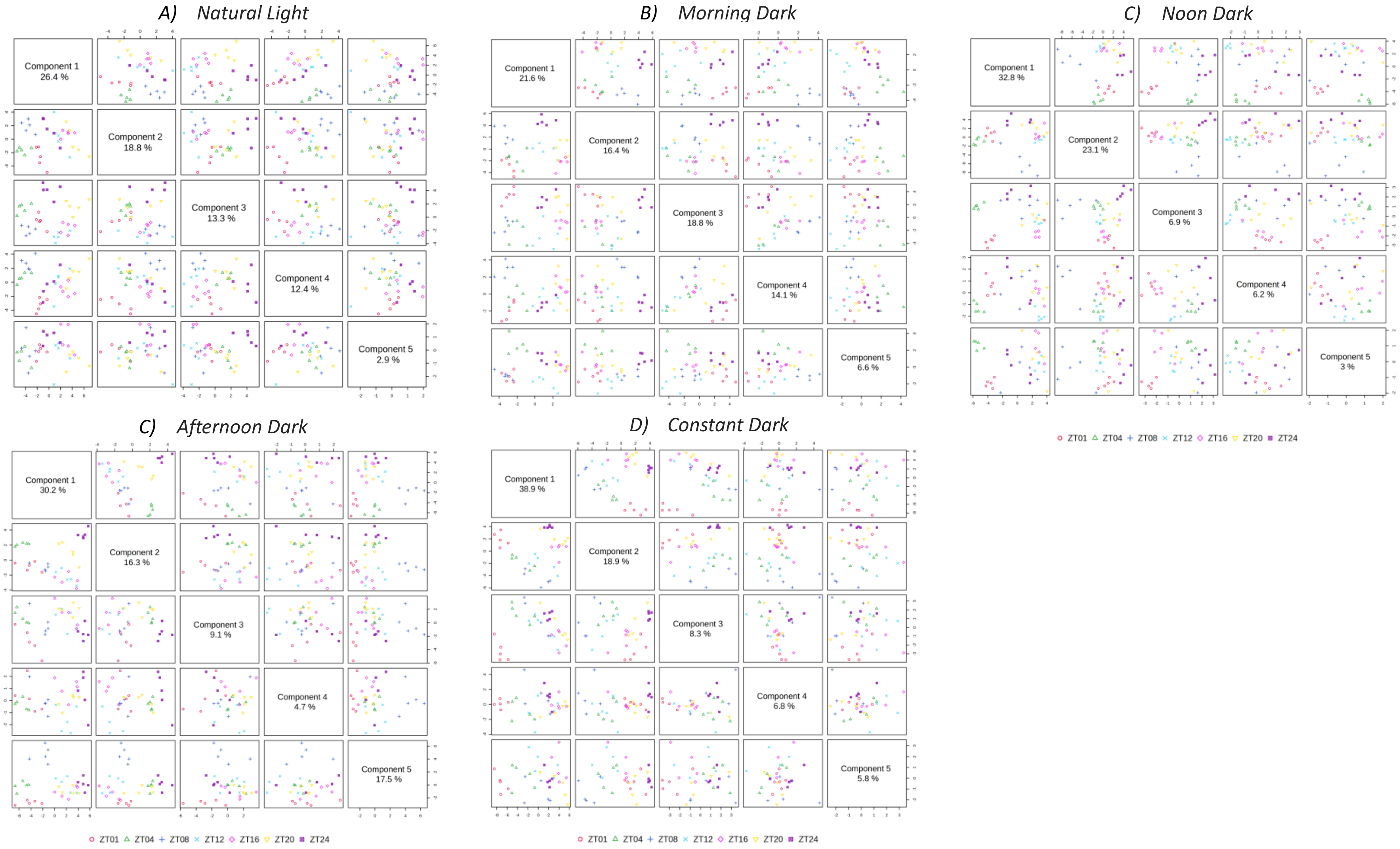


**Fig. S4. Pairwise score plots of the latent components generated by the partial least squares-discriminant analysis (PLS-DA) model for sugarcane leaves subjected to five darkness treatments throughout the diel cycle.** Each panel represents one treatment: (A) Natural Light, (B) Morning Dark, (C) Noon Dark, (D) Afternoon Dark, and (E) Constant Dark. Pairwise comparisons among the first five latent components are shown, with the percentage of variance explained by each component indicated along the diagonal. Samples correspond to different Zeitgeber times (ZTs; hours after dawn).

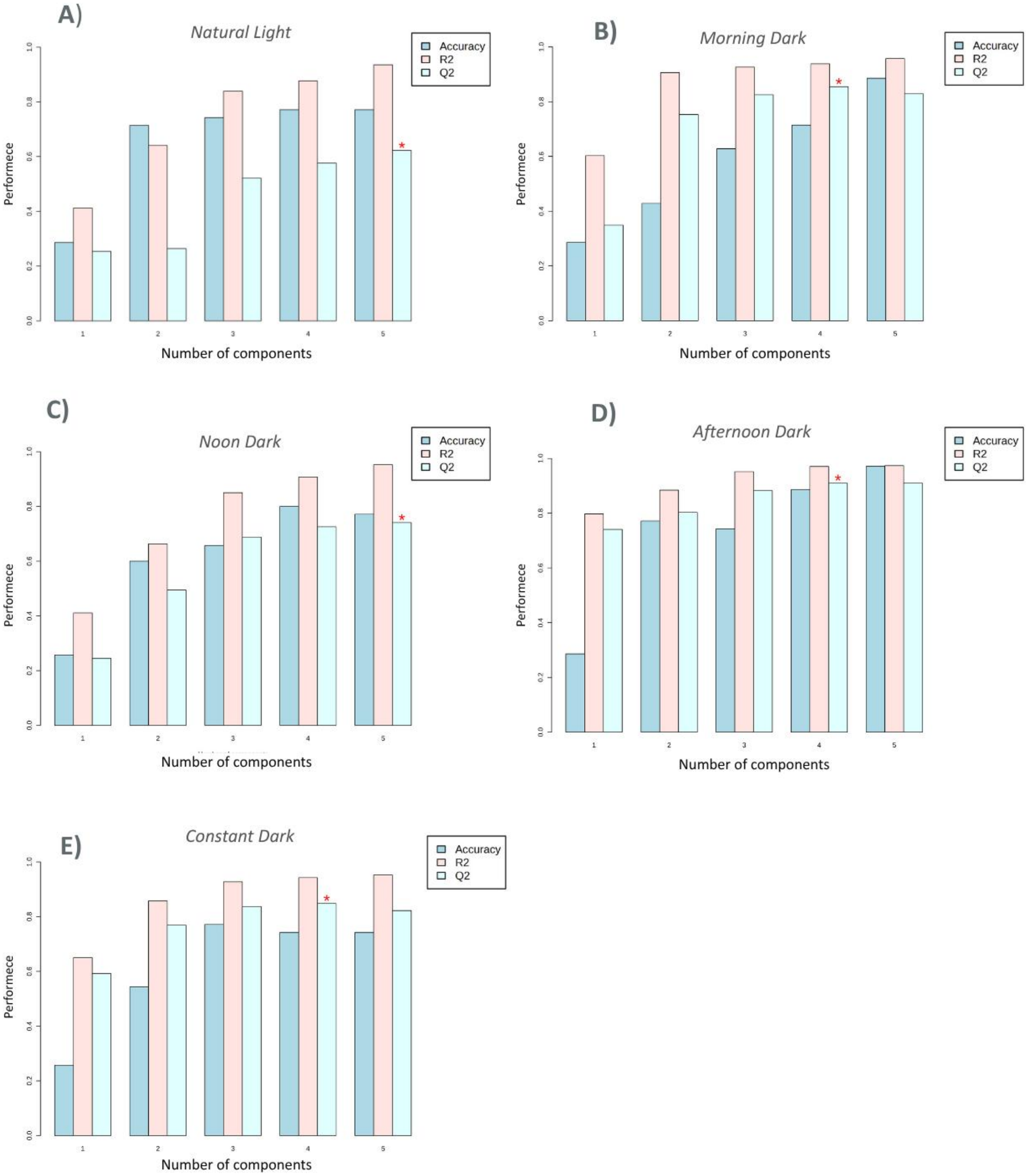


**Fig. S5. Five-fold cross-validation of the partial least squares discriminant analysis (PLS-DA) models for sugarcane leaves subjected to five darkness treatments throughout the diel cycle.** Model performance across the first five latent components is presented for each treatment: (A) Natural Light, (B) Morning Dark, (C) Noon Dark, (D) Afternoon Dark, and (E) Constant Dark. Accuracy, $R^2$, and $Q^2$ values are shown for each component to assess the predictive performance and robustness of the PLS-DA models.

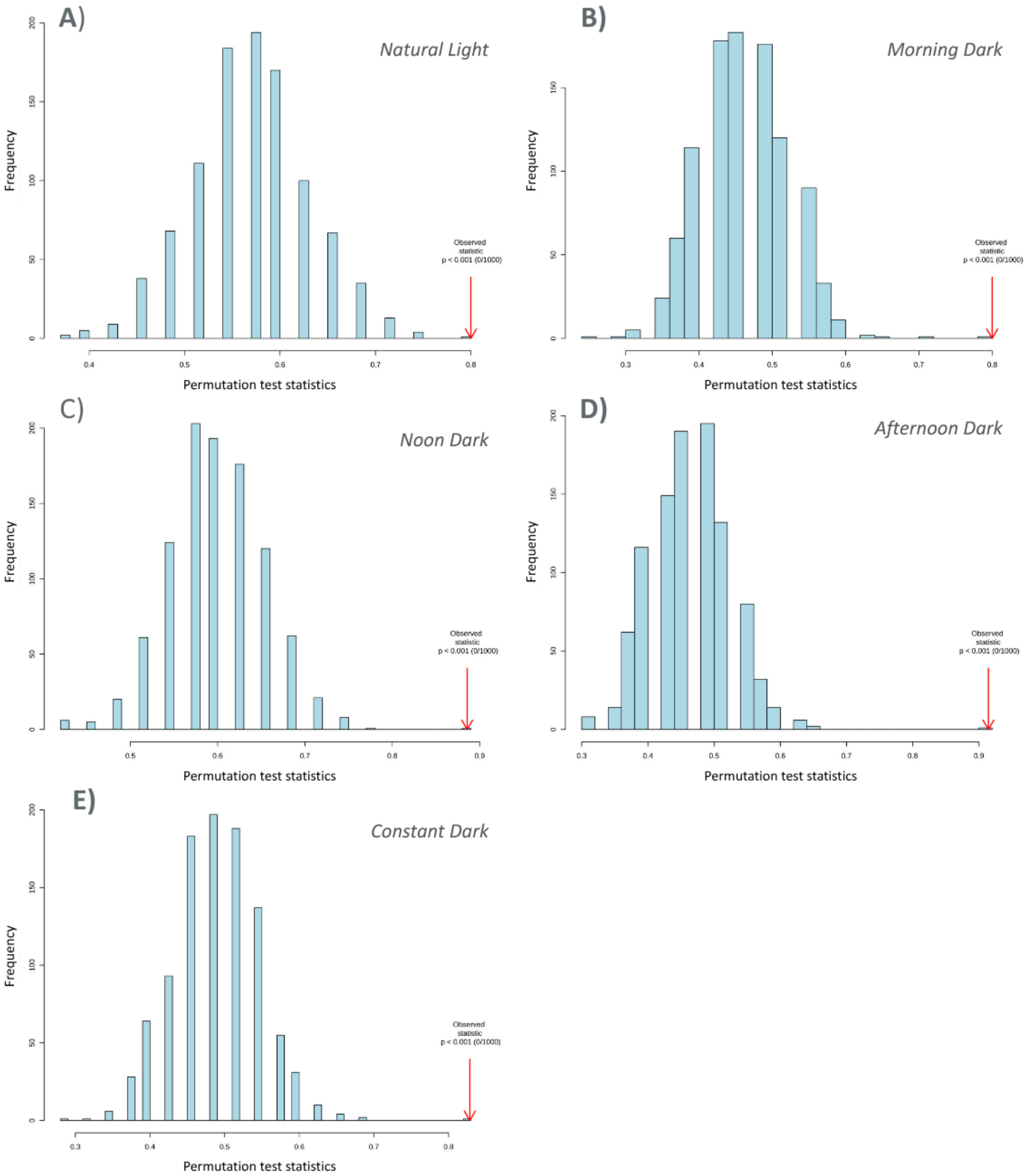


**Fig. S6. Permutation tests of the partial least squares discriminant analysis (PLS-DA) models for sugarcane leaves subjected to five darkness treatments throughout the diel cycle.** Each panel corresponds to one treatment: (A) Natural Light, (B) Morning Dark, (C) Noon Dark, (D) Afternoon Dark, and (E) Constant Dark. Model significance was assessed using prediction accuracy obtained from 1,000 permutations. The observed model performance is compared with the distribution of permuted models to evaluate whether the discrimination exceeds that expected by chance.

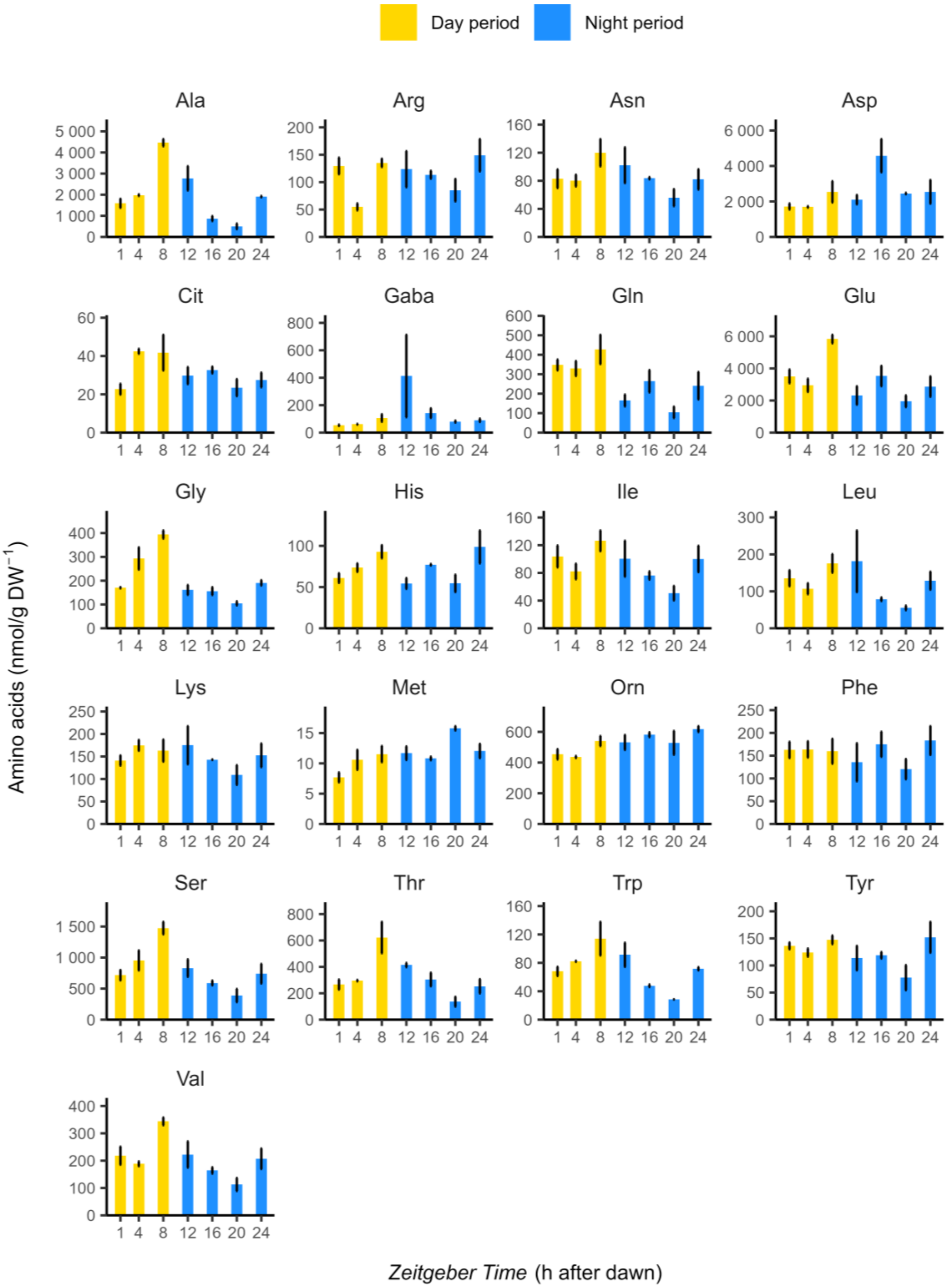


**Fig. S7. Natural Light treatment: Amino acid variation profile across the diel cycle in sugarcane leaves.** Data are presented as means ± SE (n = 5). Abbreviations: Asp, aspartate; Glu, glutamate; Asn, asparagine; Ser, serine; Gln, glutamine; His, histidine; Gly, glycine; Arg, arginine; Thr, threonine; Ala, alanine; Tyr, tyrosine; GABA, γ-aminobutyric acid; Met, methionine; Trp, tryptophan; Val, valine; Phe, phenylalanine; Ile, isoleucine; Leu, leucine; Orn, ornithine; Lys, lysine; and Cit, citrulline.

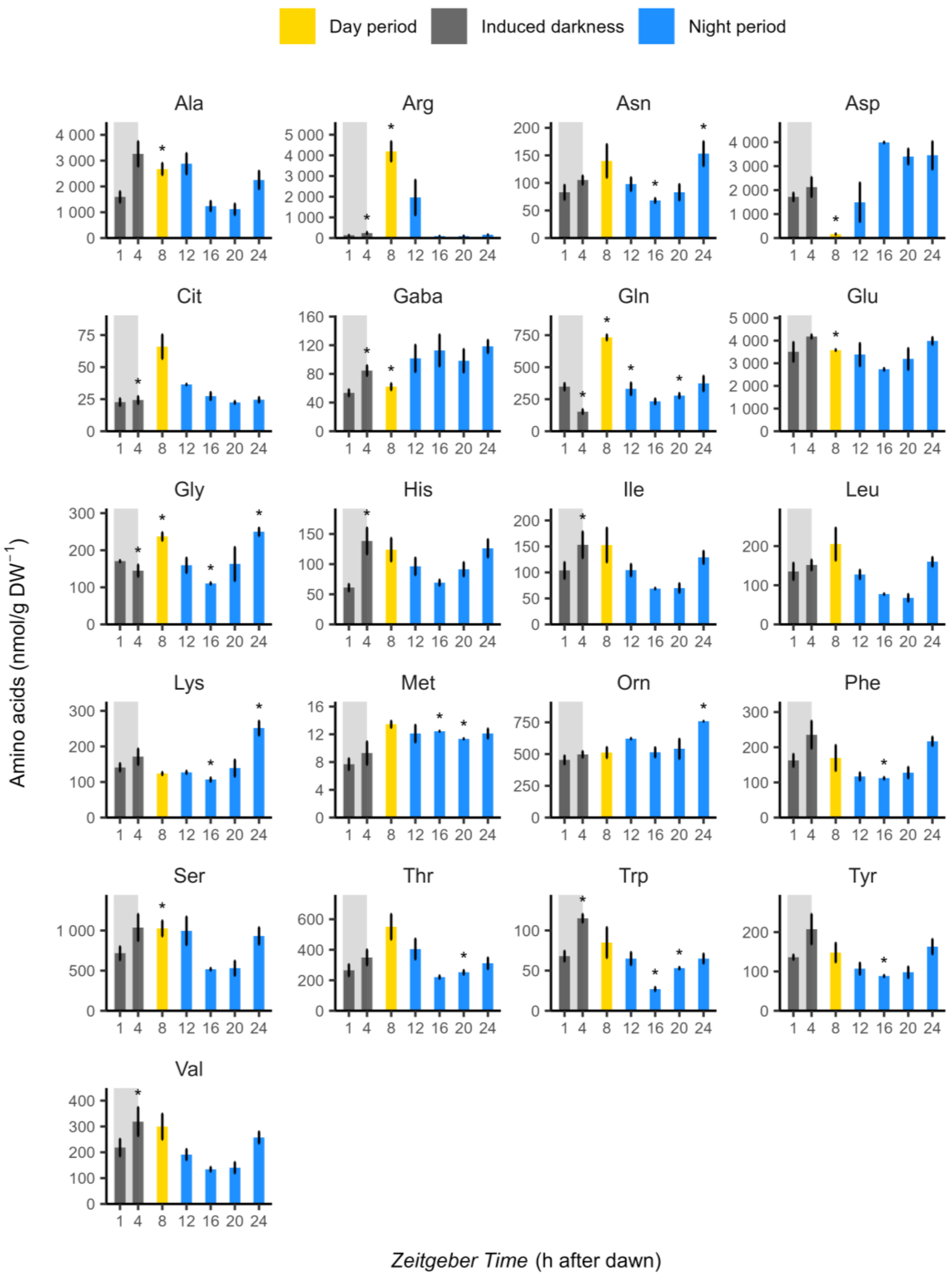


**Fig. S8 Morning dark treatment: Amino acid variation profile across the diel cycle in sugarcane leaves.** Data are presented as means ± SE (n = 5). Abbreviations: Asp, aspartate; Glu, glutamate; Asn, asparagine; Ser, serine; Gln, glutamine; His, histidine; Gly, glycine; Arg, arginine; Thr, threonine; Ala, alanine; Tyr, tyrosine; GABA, γ-aminobutyric acid; Met, methionine; Trp, tryptophan; Val, valine; Phe, phenylalanine; Ile, isoleucine; Leu, leucine; Orn, ornithine; Lys, lysine; and Cit, citrulline.

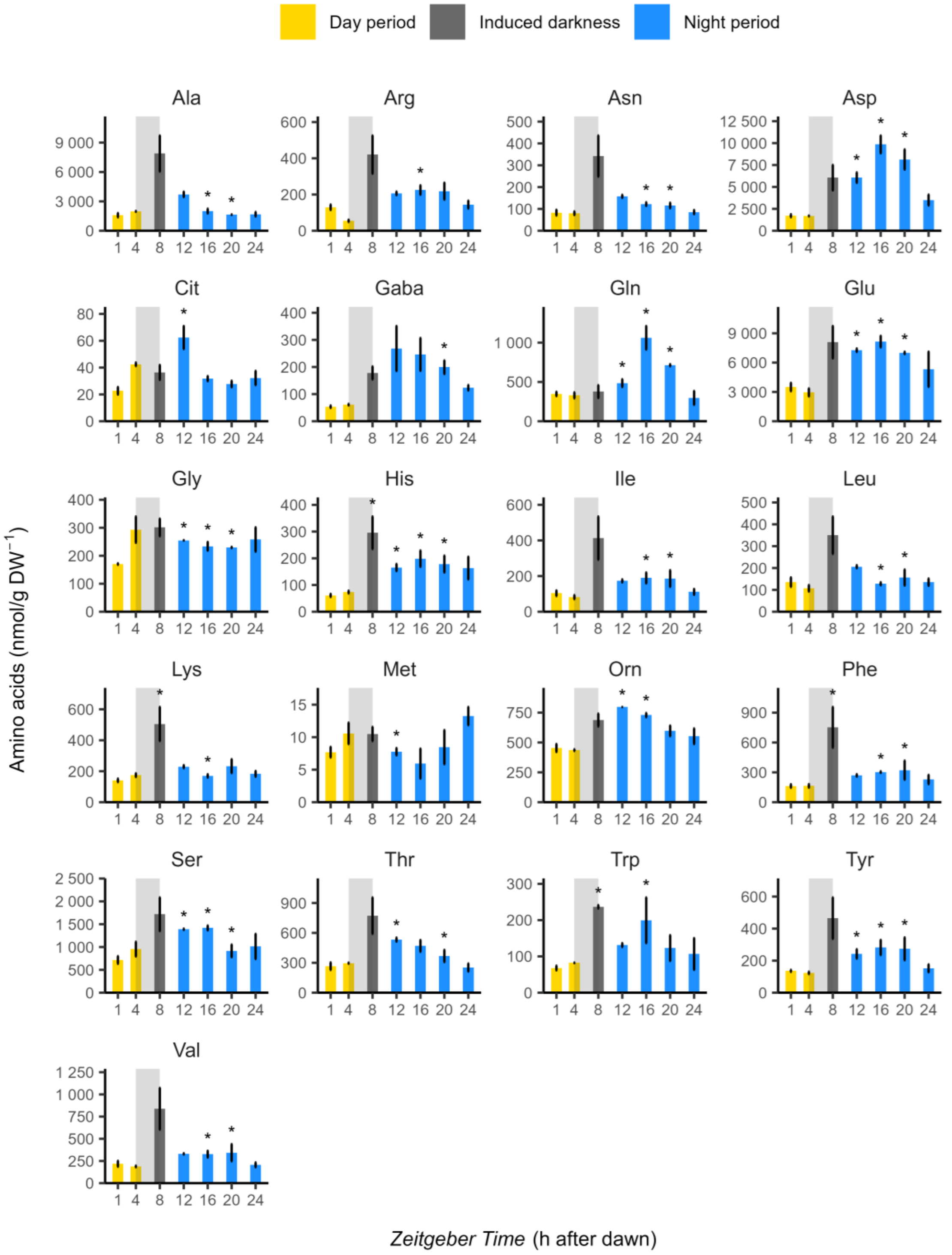


**Fig. S9. Noon dark treatment: Amino acid variation profile across the diel cycle in sugarcane leaves.** Data are presented as means ± SE (n = 5). Abbreviations: Asp, aspartate; Glu, glutamate; Asn, asparagine; Ser, serine; Gln, glutamine; His, histidine; Gly, glycine; Arg, arginine; Thr, threonine; Ala, alanine; Tyr, tyrosine; GABA, γ-aminobutyric acid; Met, methionine; Trp, tryptophan; Val, valine; Phe, phenylalanine; Ile, isoleucine; Leu, leucine; Orn, ornithine; Lys, lysine; and Cit, citrulline.

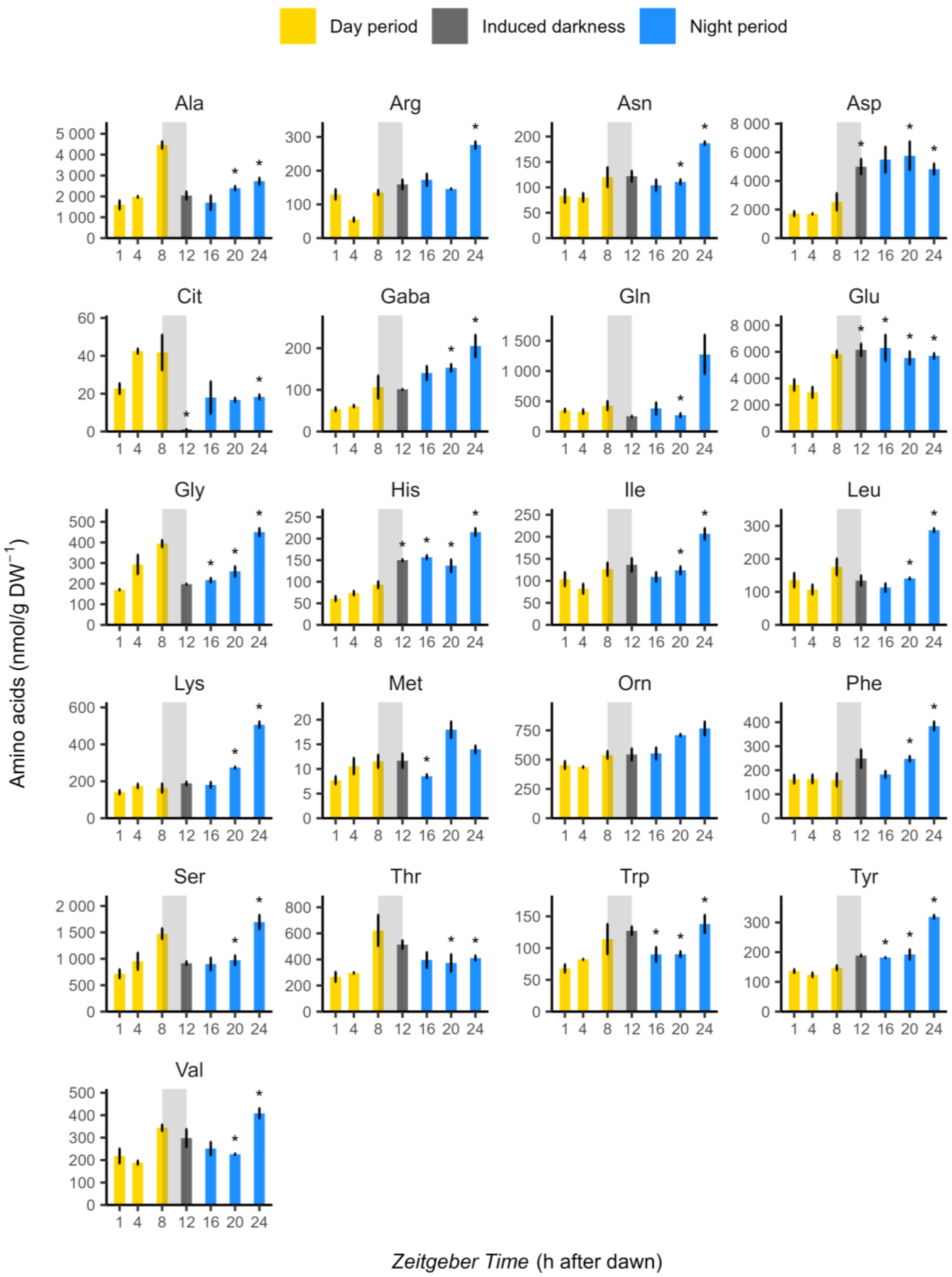


**Fig. S10. Afternoon dark treatment: Amino acid variation profile across the diel cycle in sugarcane leaves.** Data are presented as means ± SE (n = 5). Abbreviations: Asp, aspartate; Glu, glutamate; Asn, asparagine; Ser, serine; Gln, glutamine; His, histidine; Gly, glycine; Arg, arginine; Thr, threonine; Ala, alanine; Tyr, tyrosine; GABA, γ-aminobutyric acid; Met, methionine; Trp, tryptophan; Val, valine; Phe, phenylalanine; Ile, isoleucine; Leu, leucine; Orn, ornithine; Lys, lysine; and Cit, citrulline.

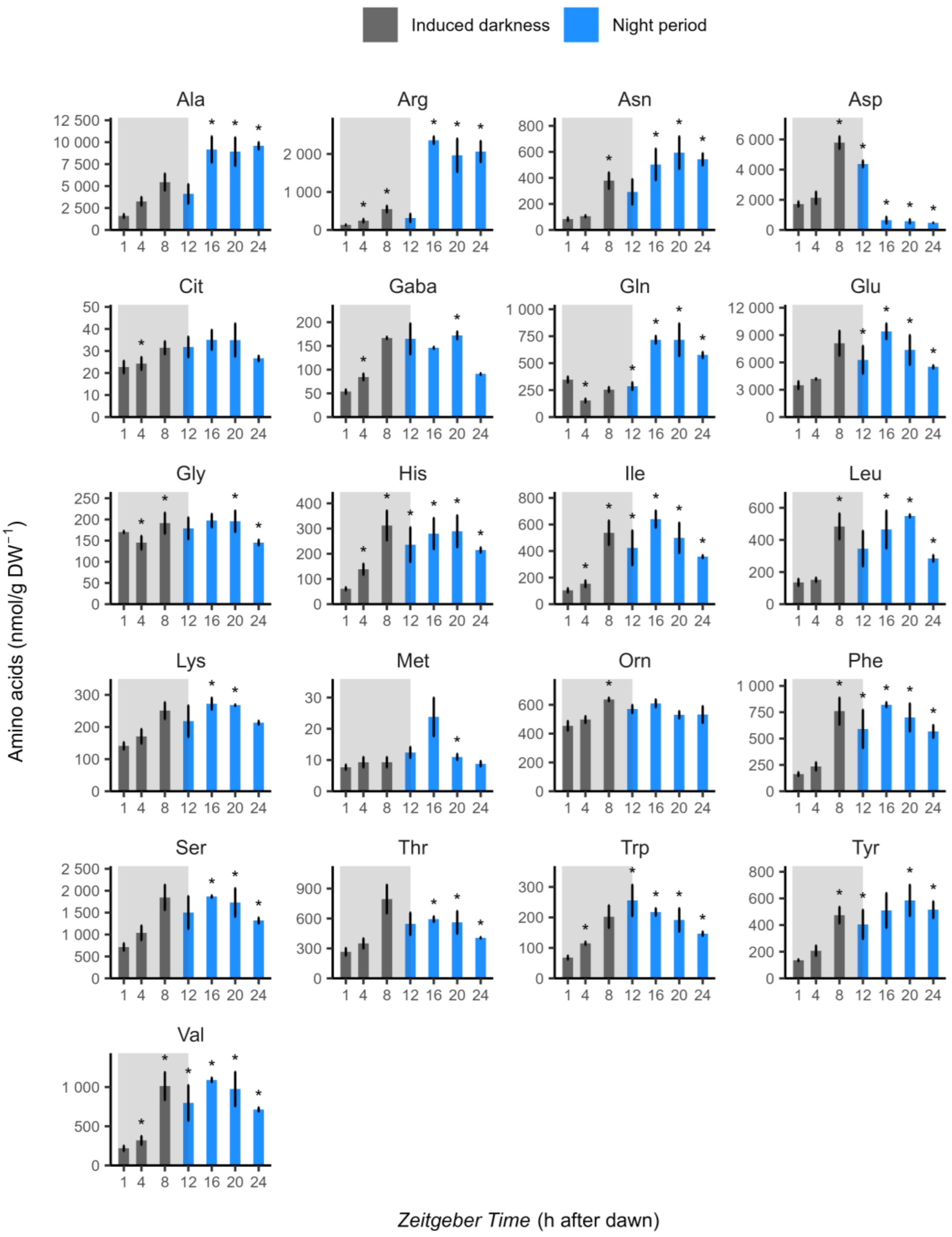


**Fig. S11. Constant dark treatment: Amino acid variation profile across the diel cycle in sugarcane leaves.** Data are presented as means ± SE (n = 5). Abbreviations: Asp, aspartate; Glu, glutamate; Asn, asparagine; Ser, serine; Gln, glutamine; His, histidine; Gly, glycine; Arg, arginine; Thr, threonine; Ala, alanine; Tyr, tyrosine; GABA, γ-aminobutyric acid; Met, methionine; Trp, tryptophan; Val, valine; Phe, phenylalanine; Ile, isoleucine; Leu, leucine; Orn, ornithine; Lys, lysine; and Cit, citrulline.

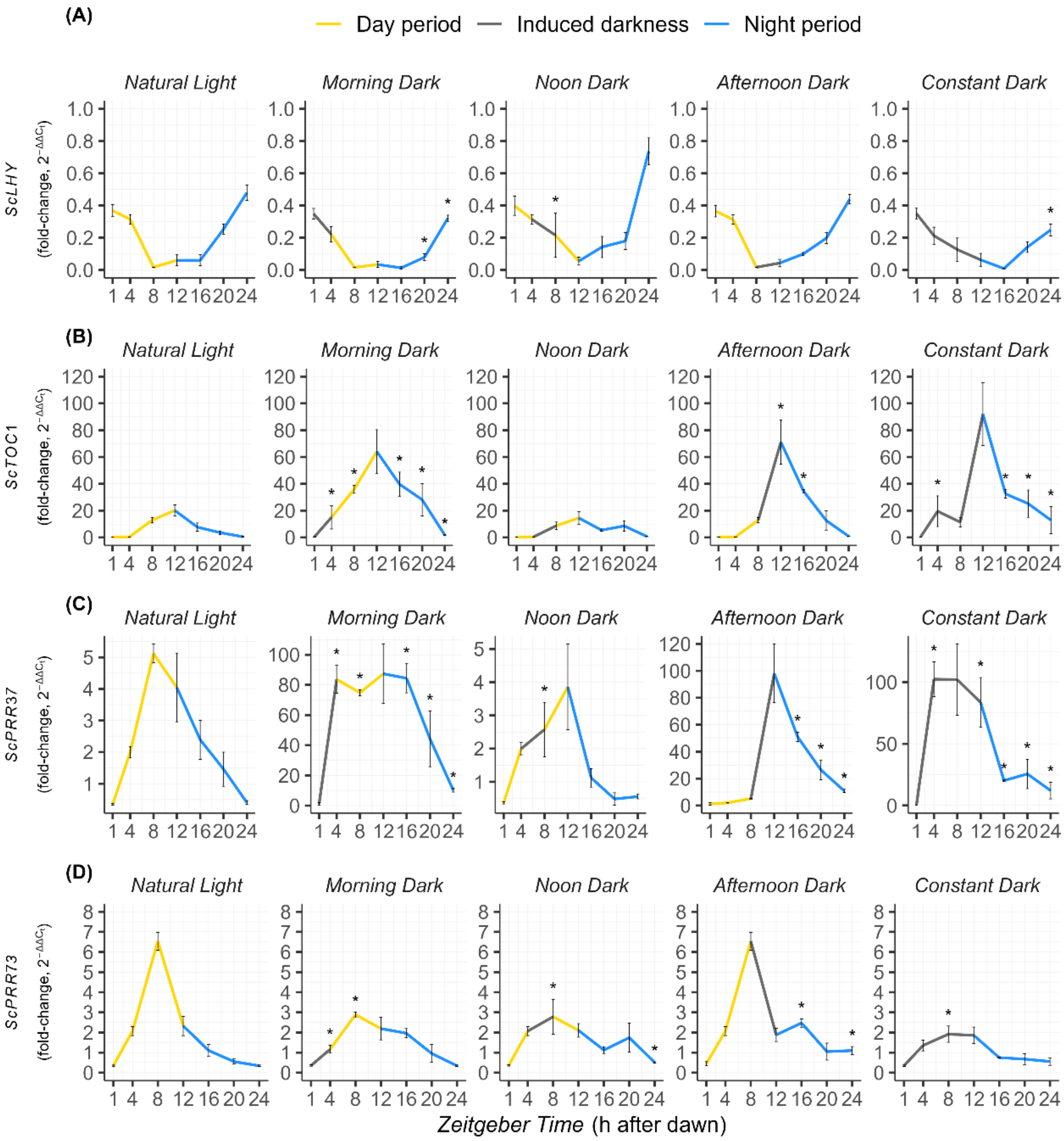


**Fig. S12. Expression profiles of core circadian clock genes in sugarcane leaves across the diel cycle under different induced-darkness treatments.** (A) *ScLHY*, (B) *ScTOC1*, (C) *ScPRR37*, and (D) *ScPRR73*. Relative transcript levels were calculated as fold-change relative to the first collection point (ZT1) and normalized to the geometric mean of the reference genes *ScGAPDH*, *ScACT*, and *ScPGR*. Yellow lines indicate the natural light period, blue lines the natural dark period (night), and grey lines the imposed darkness on leaf +1 during the photoperiod. Data are presented as means ± SE (n = 3). Statistical significance was determined using the Wilcoxon test, with asterisks indicating differences between Natural Light and dark treatments ($P < 0.05$).

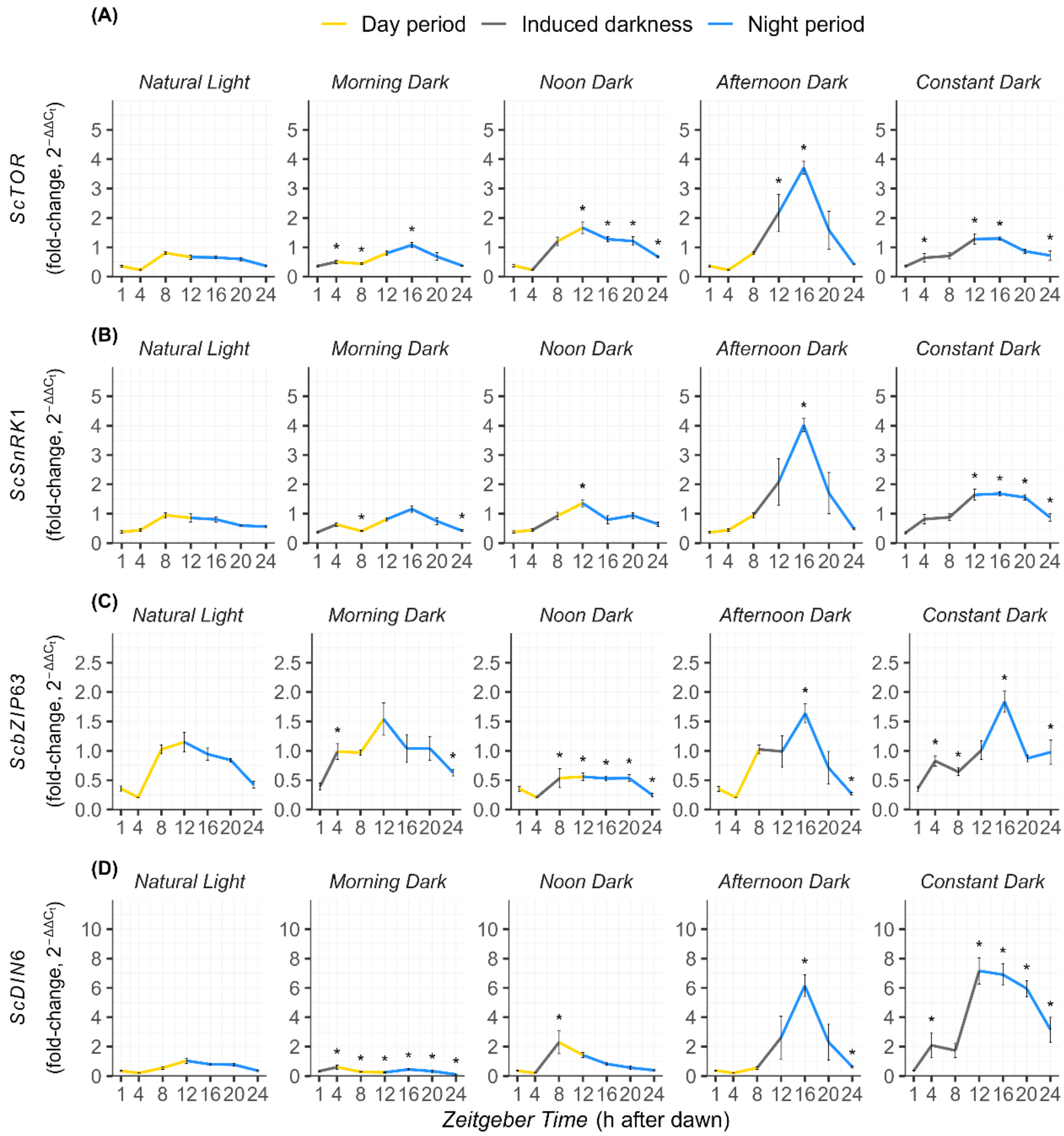


**Fig. S13. Expression profiles of sugar-sensing players and downstream targets in sugarcane leaves across the diel cycle under different induced-darkness treatments.** (A) *ScTOR*, (B) *ScSnRK1*, (C) *ScBzip63*, and (D) *ScDIN6*. Relative transcript levels were calculated as fold-change relative to the first collection point (ZT1) and normalized to the geometric mean of the reference genes *ScGAPDH*, *ScACT*, and *ScPGR*. Data are presented as means ± SE ($n = 3$). Yellow lines indicate the natural light period, blue lines the natural dark period (night) and grey lines the imposed darkness on leaf +1 during the photoperiod. Statistical significance was determined using the Wilcoxon test, with asterisks indicating differences between Natural Light and dark treatments ($P < 0.05$).

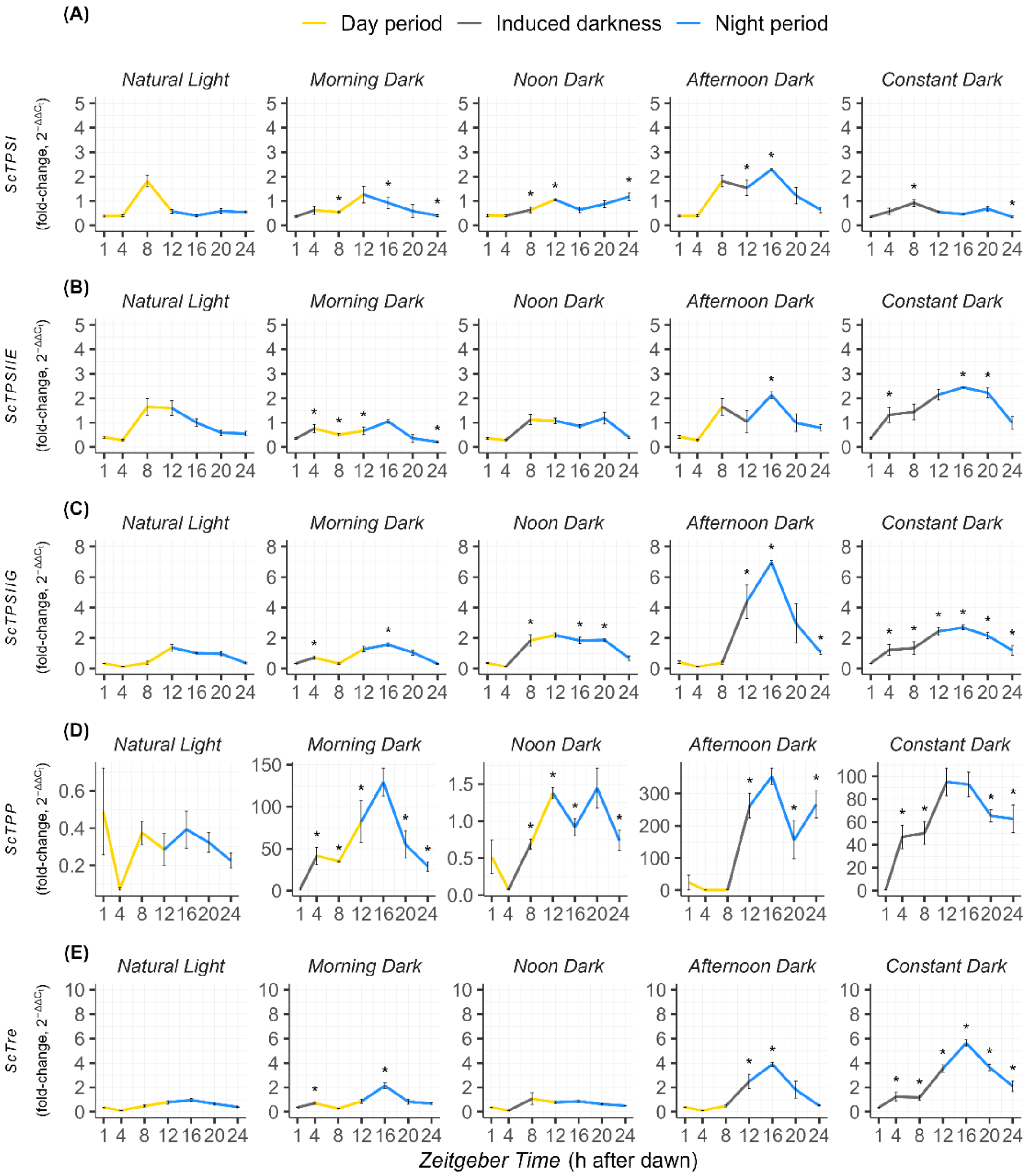


**Fig. S14. Expression profiles of trehalose pathway targets in sugarcane leaves across the diel cycle under different induced-darkness treatments.** (A) *ScTPSI*, (B) *ScTPSIIE*, (C) *ScTPSIIG*, and (D) *ScTPP* and (E*) ScTre*. Relative transcript levels were calculated as fold-change relative to the first collection point (ZT1) and normalized to the geometric mean of the reference genes *ScGAPDH*, *ScACT*, and *ScPGR*. Yellow lines indicate the natural light period, blue lines the natural dark period (night) and grey lines the imposed darkness on leaf +1 during the photoperiod. Data are presented as means ± SE (n = 3). Statistical significance was determined using the Wilcoxon test, with asterisks indicating differences between Natural Light and dark treatments (P < 0.05

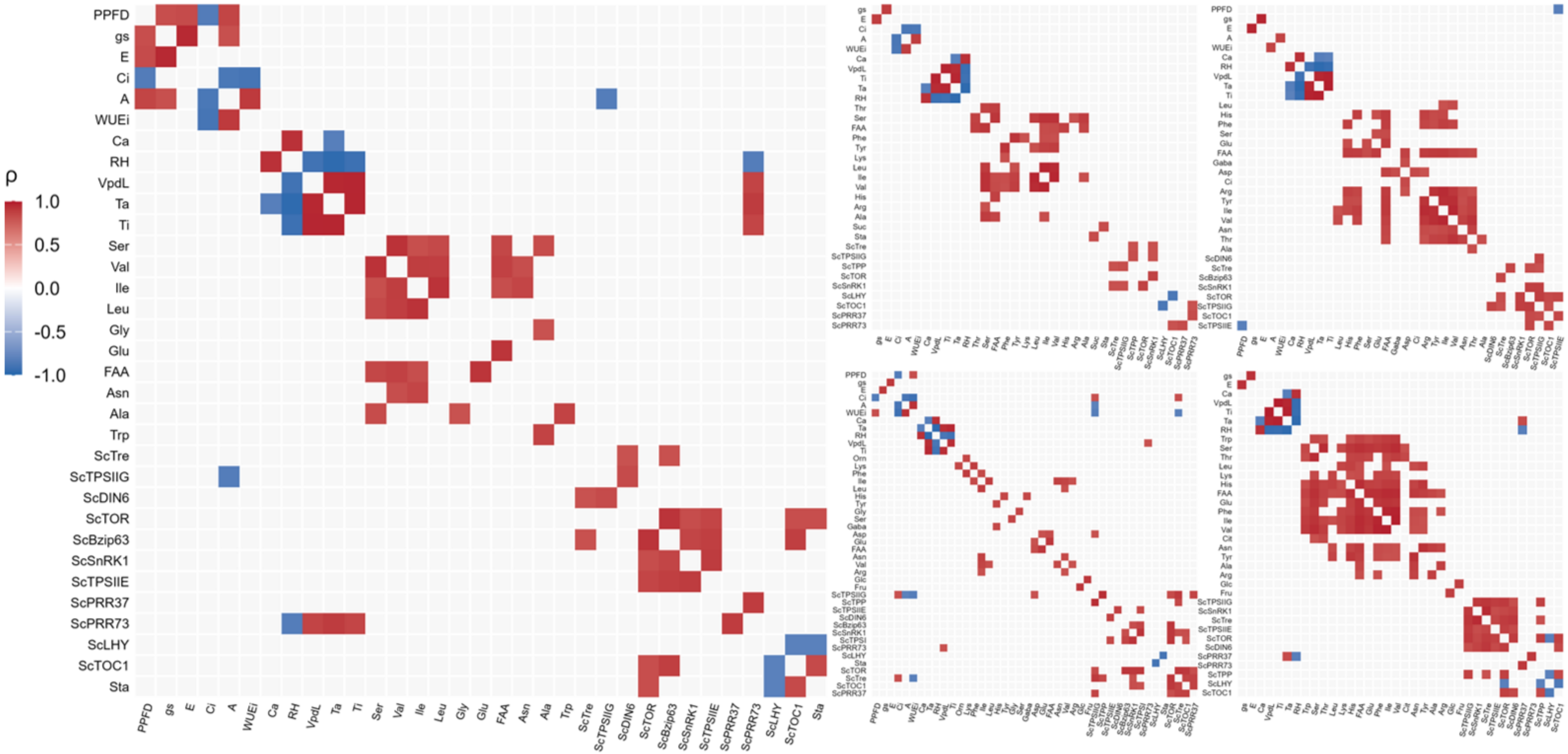


**Fig. S15. Spearman's correlation heatmaps showing the co-variation among gas-exchange parameters, non-structural carbohydrates, free amino acids, and gene expression levels in sugarcane leaves subjected to five darkness treatments:** (A) Natural Light, (B) Morning Dark, (C) Noon Dark, (D) Afternoon Dark, and (E) Constant Dark. Heatmaps display significant pairwise correlations (|r| > 0.8, $P < 0.05$; n = 5). Positive and negative correlations are represented in red and blue, respectively. Only variables with at least one significant correlation are shown in each panel. Abbreviations: A, net $CO_2$ assimilation rate; Ala, alanine; Arg, arginine; Asn, asparagine; Asp, aspartate; Ca, ambient $CO_2$ concentration; Ci, intercellular $CO_2$ concentration; Cit, citrulline; E, transpiration rate; FAA, free amino acids; Fru, fructose; GABA, γ-aminobutyric acid; Glc, glucose; Glu, glutamate; Gly, glycine; gs, stomatal conductance; His, histidine; Ile, isoleucine; Leu, leucine; Lys, lysine; Orn, ornithine; Phe, phenylalanine; PPFD, photosynthetic photon flux density; RH, relative humidity; Ser, serine; Sta, starch; Suc, sucrose; Ta, air temperature; Thr, threonine; Ti, leaf temperature; Trp, tryptophan; Tyr, tyrosine; Val, valine; VPDL, leaf-to-air vapor pressure deficit; and WUEi, intrinsic water-use efficiency.
