## Supplementary Tables for "Timing of transient darkness shapes carbon–nitrogen metabolism and sugar signaling in sugarcane"

**Table S1. List of primers used for RT-qPCR analysis. Forward (F) and reverse (R) primer sequences are shown in the 5′ to 3′ direction.**

| | | |
|---|---|---|
| *ACT* | Foward: | CCAGTTCCATTGTCACAAACAAG |
| | Reverse: | TCTCCGGAATCCGTAGCAAA |
| *PGR* | Foward: | GTTGCCGGTCATCCAGAACA |
| | Reverse: | GCGGCTTTGTCAGGGACATT |
| *UBQ10* | Foward: | CGTCCGCAGTCCCCAAT |
| | Reverse: | TGAGAGGATCGCGAGGATTC |
| *TOR* | Foward: | AGTACCATTCCGTTTGACTAGA |
| | Reverse: | TGTCTCTGTTTGTGCGAAGG |
| *SnRK1* | Foward: | CATCTCCCACGCTATTTGACT |
| | Reverse: | AGCAACAGTCGCCTCATTTT |
| *TPS class I* | Foward: | GAGAAGGTGATGTTGAAAGG |
| | Reverse: | CTGGGATTTGGAGAGGAG |
| *TPS class IIE* | Foward: | CGATGTTTCTACTATCTGAGGC |
| | Reverse: | GGACTTGTTGGGGCATAG |
| *TPS class IIG* | Foward: | CTCTATCTCTCTGAACTCTTTCC |
| | Reverse: | CTGGCTTGTGGTCAAATG |
| *TPP* | Foward: | CTCTTCCTGTTCAGCCTATTG |
| | Reverse: | GTGTTCCCTTCTTGCCATC |
| *Ter* | Foward: | CAATAAACACAAACCCACGAAC |
| | Reverse: | CAGAGGACAAGGAAATAGC |
| *DIN6* | Foward: | GGCTACCTCTACTTCCAC |
| | Reverse: | ATTGATGAACTCCTTGTCCA |
| *bZIP63* | Foward: | GATTAGCAATGTCCACTAC |
| | Reverse: | ATTTTGTCAACGGGTCTG |

**Table S2. q-values obtained from BioNetStat differential network analyses comparing the Natural Light control with each darkness treatment using Spearman's rank correlation (|r| > 0.8).** Network comparisons were performed using four node centrality metrics: degree, betweenness, closeness, and eigenvector centrality. Analyses were conducted using all variables together or grouped into seven functional sets: All genes, Sugar sensing, Circadian clock, All metabolites, Non-structural carbohydrates (NSC), Amino acids, and Gas exchange. Statistical significance was assessed by permutation testing (1,000 permutations). Dashes (–) indicate missing values resulting from network inference failures**.**

| | Morning Dark | Noon Dark | Afternoon Dark | Constant Dark |
|---|---|---|---|---|
| **Dregree centrality** | | | | |
| All variables | 0.70 | 0.23 | 0.60 | 0.03* |
| All genes | 0.18 | 0.25 | 0.04 * | 0.01 * |
| Sugar sensing | 0.17 | 0.29 | 0.15 | 0.03 * |
| Circadian clock | 0.63 | 0.08 | 0.23 | 0.73 |
| All metabolites | 0.94 | 0.30 | 0.97 | 0.05 |
| NSC | 0.48 | 1.00 | 0.13 | 0.28 |
| Amino acids | 0.94 | 0.30 | 0.97 | 0.05 |
| Gas exchange | 0.11 | 0.03 * | 0.47 | 0.03 * |
| **Betweenness centrality** | | | | |
| All variables | 0.01* | 0.01* | 0.011* | 0.01* |
| All genes | 0.04* | 0.03 * | 0.05 * | 0.04* |
| Sugar sensing | 0.03* | 0.02 * | 0.001 * | 0.03* |
| Circadian clock | 0.002* | 1.00 | 1.00 | 1.00 |
| All metabolites | 0.80 | 0.04* | 0.53 | 0.40 |
| NSC | 1.00 | 1.00 | 1.00 | 1.00 |
| Amino acids | 0.79 | 0.04* | 0.53 | 0.38 |
| Gas exchange | 0.32 | 0.71 | 0.32 | 0.29 |
| **Closeness centrality** | | | | |
| All variables | 0.26 | 0.09* | 0.05 | 0.16 |
| All genes | - | - | - | - |
| Sugar sensing | - | - | - | - |
| Circadian clock | - | - | - | - |
| All metabolites | 0.75 | 0.31 | 0.01* | 0.40 |
| NSC | - | - | - | - |
| Amino acids | 0.65 | 0.26 | 0.02* | 0.31 |
| Gas exchange | 0.23 | 0.02 * | 0.34 | 0.04 * |
| **Eigenvector centrality** | | | | |
| All variables | 0.25 | 0.16 | 0.20 | 0.04* |
| All genes | 0.10 | 0.43 | 0.30 | 0.16 |
| Sugar sensing | 0.04* | 0.22 | 0.32 | 0.06 |
| Circadian clock | 0.48 | 0.45 | 0.46 | 0.70 |
| All metabolites | 0.80 | 0.03* | 0.71 | 0.03* |
| NSC | 0.43 | 1.00 | 0.45 | 0.43 |
| Amino acids | 0.78 | 0.02* | 0.73 | 0.05* |
| Gas exchange | 0.69 | 0.78 | 0.95 | 0.74 |

**Table S3. Changes in pairwise variable associations between the Natural Light control and darkness treatments estimated using BioNetStat.** Values correspond to the difference in Spearman's correlation coefficients (Δr) between each darkness treatment and the Natural Light condition for variable pairs with significant correlations (|r| > 0.8). Positive and negative values indicate increases and decreases in association strength, respectively.

| | | Morning Dark | Noon Dark | Afternoon Dark | Constant Dark |
|---|---|---|---|---|---|
| **Gas-exchange** | | | | | |
| *A* | $C_i$ | 0.03 | -0.87 | 0.01 | -0.87 |
| *A* | gs | -0.81 | -0.81 | -0.81 | -0.81 |
| *A* | PPFD | -0.86 | -0.86 | -0.86 | -0.86 |
| *A* | $WUE_i$ | 0.02 | -0.81 | 0.03 | -0.90 |
| $C_a$ | $T_i$ | - | 0.80 | - | - |
| $C_a$ | $Vpd_L$ | 0.83 | - | - | 0.08 |
| $C_i$ | PPFD | -0.84 | -0.84 | -0.83 | -0.84 |
| $C_i$ | $WUE_i$ | 0.01 | -0.89 | 0.03 | -0.06 |
| *E* | PPFD | -0.83 | -0.83 | -0.83 | -0.83 |
| gs | E | -0.09 | 0.01 | -0.06 | -0.06 |
| gs | PPFD | -0.82 | -0.82 | -0.82 | -0.82 |
| PPFD | $WUE_i$ | - | - | 0.81 | - |
| RH | $T_i$ | -0.04 | 0.02 | -0.01 | - |
| RH | $Vpd_L$ | 0.04 | - | -0.03 | 0.05 |
| $T_a$ | $T_i$ | -0.26 | 0.17 | -0.10 | -0.07 |
| $T_a$ | $Vpd_L$ | 0.06 | -8.63 | -0.21 | 0.22 |
| $Vpd_L$ | $T_i$ | -0.10 | 0.09 | -0.28 | -0.07 |
| **Sugars** | | | | | |
| Glc | Fru | - | - | 0.91 | 0.87 |
| Suc | Sta | 0.83 | - | - | - |
| **Amino acids** | | | | | |
| Ala | Arg | - | - | - | 0.87 |
| Ala | Asn | - | - | - | 0.88 |
| Ala | Gly | -0.80 | -0.80 | -0.80 | -0.80 |
| Ala | Ile | 0.81 | - | - | - |
| Ala | Ser | 0.06 | -0.83 | -0.83 | -0.83 |
| Ala | Thr | - | 0.82 | - | - |
| Ala | Trp | -0.86 | -0.86 | -0.86 | -0.86 |
| Arg | Asn | - | 0.83 | - | 0.09 |
| Arg | Asp | - | 0.80 | - | - |
| Arg | His | - | 0.87 | - | - |
| Arg | Ile | - | 0.09 | 0.80 | - |
| Arg | Phe | - | 0.85 | - | 0.80 |
| Arg | Ser | 0.80 | - | - | - |
| Arg | Thr | - | 0.81 | - | - |
| Arg | Tyr | - | 0.92 | - | - |
| Arg | Val | - | 0.91 | 0.80 | - |

| | | | | | |
|---|---|---|---|---|---|
| Asn | His | - | - | - | 0.09 |
| Asn | Ile | -0.86 | -0.02 | 0.01 | -0.04 |
| Asn | Leu | - | - | - | 0.83 |
| Asn | Phe | - | - | - | 0.82 |
| Asn | Thr | - | 0.87 | - | - |
| Asn | Tyr | - | 0.82 | - | 0.93 |
| Asn | Val | -0.82 | 0.05 | 0.02 | -0.01 |
| Asp | $C_i$ | - | 0.84 | - | - |
| Asp | Gaba | - | 0.82 | - | - |
| Asp | Glu | - | - | 0.80 | - |
| Cit | Ser | - | - | - | 0.80 |
| Cit | Thr | - | - | - | 0.84 |
| Gaba | His | - | - | 0.84 | - |
| Glu | His | - | 0.82 | - | 0.87 |
| Glu | Ile | - | - | - | 0.92 |
| Glu | Phe | - | - | - | 0.90 |
| Glu | Ser | - | 0.08 | - | 0.91 |
| Glu | Thr | - | - | - | 0.85 |
| Glu | Trp | - | - | - | 0.84 |
| Glu | Val | - | - | - | 0.91 |
| Gly | Ser | - | - | 0.85 | - |
| His | Ile | - | 0.84 | - | 0.89 |
| His | Leu | - | - | - | 0.87 |
| His | Lys | - | - | - | 0.81 |
| His | Phe | - | 0.88 | - | 0.87 |
| His | Ser | - | - | - | 0.91 |
| His | Thr | - | - | - | 0.86 |
| His | Trp | - | - | - | 0.82 |
| His | Tyr | - | 0.83 | 0.84 | 0.90 |
| His | Val | - | - | - | 0.90 |
| Ile | Leu | 0.04 | -0.12 | -0.83 | -0.11 |
| Ile | Lys | - | - | - | 0.86 |
| Ile | Phe | 0.82 | 0.88 | 0.83 | 0.97 |
| Ile | Ser | 0.11 | -0.82 | -0.82 | 0.11 |
| Ile | Thr | - | 0.87 | - | 0.85 |
| Ile | Trp | - | - | - | 0.89 |
| Ile | Tyr | 0.87 | 0.93 | - | 0.80 |
| Ile | Val | 0.08 | 0.07 | -0.03 | 0.10 |
| Leu | Lys | - | - | - | 0.85 |
| Leu | Ser | 0.05 | -0.84 | -0.84 | -0.84 |
| Leu | Tyr | 0.81 | - | - | 0.85 |
| Leu | Val | -0.79 | -0.05 | -0.08 | -0.88 |
| Lys | Orn | - | - | 0.84 | - |
| Lys | Phe | 0.80 | - | 0.09 | 0.85 |
| Lys | Ser | - | - | - | 0.84 |
| Lys | Val | - | - | - | 0.85 |

| | | | | | |
|---|---|---|---|---|---|
| Phe | Ser | - | - | - | 0.90 |
| Phe | Trp | - | - | - | 0.87 |
| Phe | Tyr | 0.09 | 0.83 | - | 0.82 |
| Phe | Val | 0.81 | 0.85 | - | 0.97 |
| Ser | Thr | 0.87 | - | - | 0.94 |
| Ser | Trp | - | - | - | 0.84 |
| Ser | Val | - | -0.93 | -0.93 | 0.02 |
| Thr | Trp | - | - | - | 0.82 |
| Thr | Val | - | 0.92 | - | 0.88 |
| Trp | Val | - | - | - | 0.90 |
| Tyr | Thr | - | 0.87 | - | - |
| Tyr | Val | 0.88 | 0.09 | - | 0.81 |
| **Cicardian clock genes** | | | | | |
| *ScLHY* | *ScTOC1* | 0.06 | -0.81 | -0.81 | 0.05 |
| *ScLHY* | *ScTOR* | - | - | - | 0.81 |
| *ScLHY* | *ScTPP* | - | - | - | 0.81 |
| *ScLHY* | Sta | -0.81 | -0.81 | 0.08 | -0.81 |
| *ScPRR37* | RH | - | - | - | 0.82 |
| *ScPRR37* | *ScPRR73* | -0.06 | -0.89 | -0.89 | 0.01 |
| *ScPRR37* | *ScTOR* | - | - | 0.81 | - |
| *ScPRR37* | *ScTPSIIG* | - | - | 0.85 | - |
| *ScPRR37* | *ScTre* | - | - | 0.87 | - |
| *ScPRR37* | $T_a$ | - | - | - | 0.83 |
| *ScPRR37* | $Vpd_L$ | - | - | - | 0.80 |
| *ScPRR73* | RH | -0.08 | -0.08 | -0.08 | -0.08 |
| *ScPRR73* | $T_a$ | -0.89 | -0.89 | -0.89 | -0.89 |
| *ScPRR73* | $T_i$ | -0.09 | -0.09 | -0.09 | -0.09 |
| *ScPRR73* | $Vpd_L$ | -0.86 | -0.86 | -0.06 | -0.86 |
| *ScTOC1* | *ScBzip63* | -0.88 | -0.88 | -0.88 | -0.88 |
| *ScTOC1* | *ScDIN6* | - | - | - | 0.84 |
| *ScTOC1* | *ScPRR37* | - | - | 0.87 | - |
| *ScTOC1* | *ScPRR73* | 0.80 | - | - | - |
| *ScTOC1* | *ScSnRK1* | - | - | 0.82 | - |
| *ScTOC1* | *ScTOR* | -0.82 | 0.02 | -0.73 | 0.06 |
| *ScTOC1* | *ScTPP* | - | - | - | 0.84 |
| *ScTOC1* | *ScTPSI* | - | - | 0.80 | - |
| *ScTOC1* | *ScTPSIIE* | - | 0.85 | - | - |
| *ScTOC1* | *ScTPSIIG* | - | 0.82 | - | - |
| *ScTOC1* | *ScTre* | - | - | 0.86 | - |
| *ScTOC1* | Sta | -0.84 | -0.84 | -0.84 | -0.84 |
| **Sugar Sensing Genes** | | | | | |
| *ScSnRK1* | *ScBzip63* | -0.86 | -0.86 | -0.04 | -0.86 |
| *ScSnRK1* | *ScDIN6* | - | - | - | 0.86 |
| *ScSnRK1* | *ScTPSI* | - | - | 0.92 | - |
| *ScSnRK1* | *ScTPSIIE* | -0.89 | -0.89 | -0.89 | -0.07 |
| *ScSnRK1* | *ScTPSIIG* | 0.81 | 0.88 | - | 0.90 |

| | | | | | |
|---|---|---|---|---|---|
| *ScSnRK1* | *ScTre* | 0.81 | - | 0.08 | 0.95 |
| *ScTOR* | *ScBzip63* | -0.92 | -0.92 | -0.01 | -0.92 |
| *ScTOR* | *ScDIN6* | - | - | - | 0.89 |
| *ScTOR* | *ScSnRK1* | 0.78 | 0.79 | 0.82 | 0.76 |
| *ScTOR* | *ScTPP* | - | - | - | 0.09 |
| *ScTOR* | *ScTPSI* | - | - | 0.90 | - |
| *ScTOR* | *ScTPSIIE* | -0.85 | -0.01 | -0.85 | 0.02 |
| *ScTOR* | *ScTPSIIG* | - | 0.88 | 0.83 | 0.90 |
| *ScTOR* | *ScTre* | - | 0.08 | 0.91 | 0.89 |
| ScTOR | Sta | -0.82 | -0.82 | -0.82 | -0.82 |
| *ScTPP* | *ScTre* | 0.08 | - | 0.84 | 0.80 |
| *ScTPSI* | *ScBzip63* | - | - | 0.86 | - |
| *ScTPSI* | *ScTPSIIE* | - | - | 0.84 | - |
| *ScTPSIIE* | PPFD | - | 0.82 | - | - |
| *ScTPSIIE* | *ScBzip63* | -0.88 | -0.88 | -0.88 | -0.88 |
| *ScTPSIIE* | *ScDIN6* | - | - | 0.87 | 0.83 |
| *ScTPSIIE* | *ScTPSIIG* | - | - | - | 0.89 |
| *ScTPSIIE* | *ScTre* | - | - | - | 0.08 |
| *ScTPSIIG* | *A* | -0.08 | -0.08 | 0.72 | -0.08 |
| *ScTPSIIG* | Asp | - | - | 0.80 | - |
| *ScTPSIIG* | $C_i$ | 0.82 | - | 0.81 | - |
| *ScTPSIIG* | *ScDIN6* | -0.83 | 0.02 | -0.83 | -0.01 |
| *ScTPSIIG* | *ScTPP* | 0.82 | - | 0.94 | 0.87 |
| *ScTPSIIG* | *ScTre* | - | 0.80 | 0.90 | 0.91 |
| *ScTPSIIG* | $WUE_i$ | 0.83 | - | 0.08 | - |
| *ScTre* | $C_i$ | - | - | 0.81 | - |
| *ScTre* | *ScBzip63* | -0.08 | 0.77 | -0.08 | -0.08 |
| *ScTre* | *ScDIN6* | -0.80 | -0.80 | -0.80 | 0.10 |
| *ScTre* | $WUE_i$ | - | - | 0.82 | - |
